# Controllable All-Atom Protein Generation with Latent Diffusion

**DOI:** 10.1101/2024.12.02.626353

**Authors:** Amy X. Lu, Jason Yang, Yueming Long, Theophile Lambert, Wilson Yan, Sarah A. Robinson, Kevin K. Yang, Vladimir Gligorijevic, Kyunghyun Cho, Richard Bonneau, Nathan C. Frey, Frances Arnold, Pieter Abbeel

**Author notes:** Contributing authors.

## Abstract

**Purpose:** Designing proteins with atomic-level functional control remains a central challenge in de novo design, exacerbated by limitations in structural data availability.

**Methods:** We introduce PLAID (Protein Latent Induced Diffusion), a generative model that efficiently co-generates discrete sequence and all-atom structure by sampling directly in the shared sequence–structure latent of a pretrained sequence-to-structure predictor. Unlike existing *de novo* generative models, PLAID trains its diffusion model exclusively on sequences, expanding effective training corpora by 2–4 orders of magnitude relative to structural databases. Using classifier-free guidance, PLAID supports controllable generation based on function (Gene Ontology) and organism keywords.

**Results:** *In silico*, PLAID can unconditionally generate all-atom structures without explicit structural supervision during diffusion model training. Function conditioned proteins can recapitulates catalytic side-chain positions for residues at non-adjacent positions, and transmembrane proteins with expected hydrophobicity patterns and predicted topologies. We further experimentally validate that PLAID can be prompted to generate heme binding proteins with high sequence novelty.

**Conclusion:** Overall, PLAID unifies sequence-scale training with atomic-level generation, enabling more precise functional control in protein design. Code and weights: github.com/amyxlu/plaid.

## 1 Introduction

Designing proteins with atomic-level functional control is a central goal in *de novo* design. Many target behaviors, such as catalysis, binding, and transport, rely on side-chain functional groups attached to a compatible backbone, i.e., the all-atom structure. Computational workflows typically split all-atom generation into two parts [1]: propose a structure and then design a sequence to fold to it (inverse-folding [2–6]), or propose a sequence and predict its structure [7–9]. Structure-first approaches afford control when a function reduces to a motif, but are constrained by the size and crystallization bias of structural databases [10, 11]. Sequence-first approaches can interpolate from evolutionary-scale training data, but provide only coarse control over atomistic determinants of function.

Another axis of limitation is functional controllability. Many backbone-level diffusion and flow-matching models tackle functional control from the direction of motif scaffolding in 3D [12–17]. This reduces reliance on traditional scaffolding libraries and is useful for binder design. However, many functions cannot be clearly determined as a motif, such as enzymatic plastic degradation or other new-to-nature biocatalytic processes. Furthermore, all-atom generation often relies on separate steps for generating sequence and structure [18, 19], or generate only backbone and sequence without sidechain atom positions [20].

Inspired by recent progress in vision-language-action (VLA) models in robot learning, which leverage information in the *weights* of vision-language models (VLMs) trained on Internet-scale data [21–24], we ask if we can leverage knowledge of protein structure captured by the weights of highly capable protein folding models [9] and learn to sample from its latent space. Doing so allows us to train at the sequence database scale, thus offering rich annotations for drastically expanded axes of control. Furthermore, existing models trained on PDB-sized corpora are biased towards crystallizable proteins. In contrast, sequence corpora are 2 to 4 orders of magnitude larger, and include coverage of intrinsically disordered regions. Additionally, by training in the compressed latent space of a trained protein folding model, we can achieve dramatic sampling speed improvements, and democratize information from highly capable trained protein models.

We introduce PLAID (Protein Latent Induced Diffusion), which learns a diffusion model in a compressed latent (CHEAP) [25], then decodes with frozen heads to recover sequence and all-atom structure. PLAID can furthermore be compositionally prompted by both functional (specifically, Gene Ontology [26]) and taxonomic keywords. *In silico*, PLAID recovers active-site side-chains of enzymes, assembles membrane proteins with correct hydropathy patterns, and can be efficiently scaled due to a fully-attention based Diffusion Transformer [27] architecture that make use of accelerated attention kernels [28, 29] and trained up to 2 billion parameters. Through wet-lab validation, we also demonstrate that function-conditioned generation from PLAID enables successful design of heme binding proteins, with designs showing high novelty in sequence space and some designs having novel structural folds.

## 2 A model for all-atom co-generation from sequence-only training

Designing proteins with atomic-level accuracy traditionally requires training on structure databases, which is biased towards crystallizable regions (e.g. ignoring intrinsically disordered regions), and is inherently data limited. To overcome this, we developed PLAID (Protein Latent Induced Diffusion), a framework that performs diffusion directly within the latent space of a pretrained protein folding model (Figure 1a–d). PLAID integrates four components: (1) extraction of a joint latent representation of protein sequence and structure; (2) latent compression to stabilize and regularize the diffusion process; (3) compositional conditioning for functional and taxonomic control; and (4) a highly scalable Transformer-based architecture. Together, these elements enable end-to-end co-generation of sequence and all-atom structure with a two billion parameter model, while requiring only sequence data to train the generative model, thus augmenting data by 2–4 orders of magnitude.

**Fig. 1:**
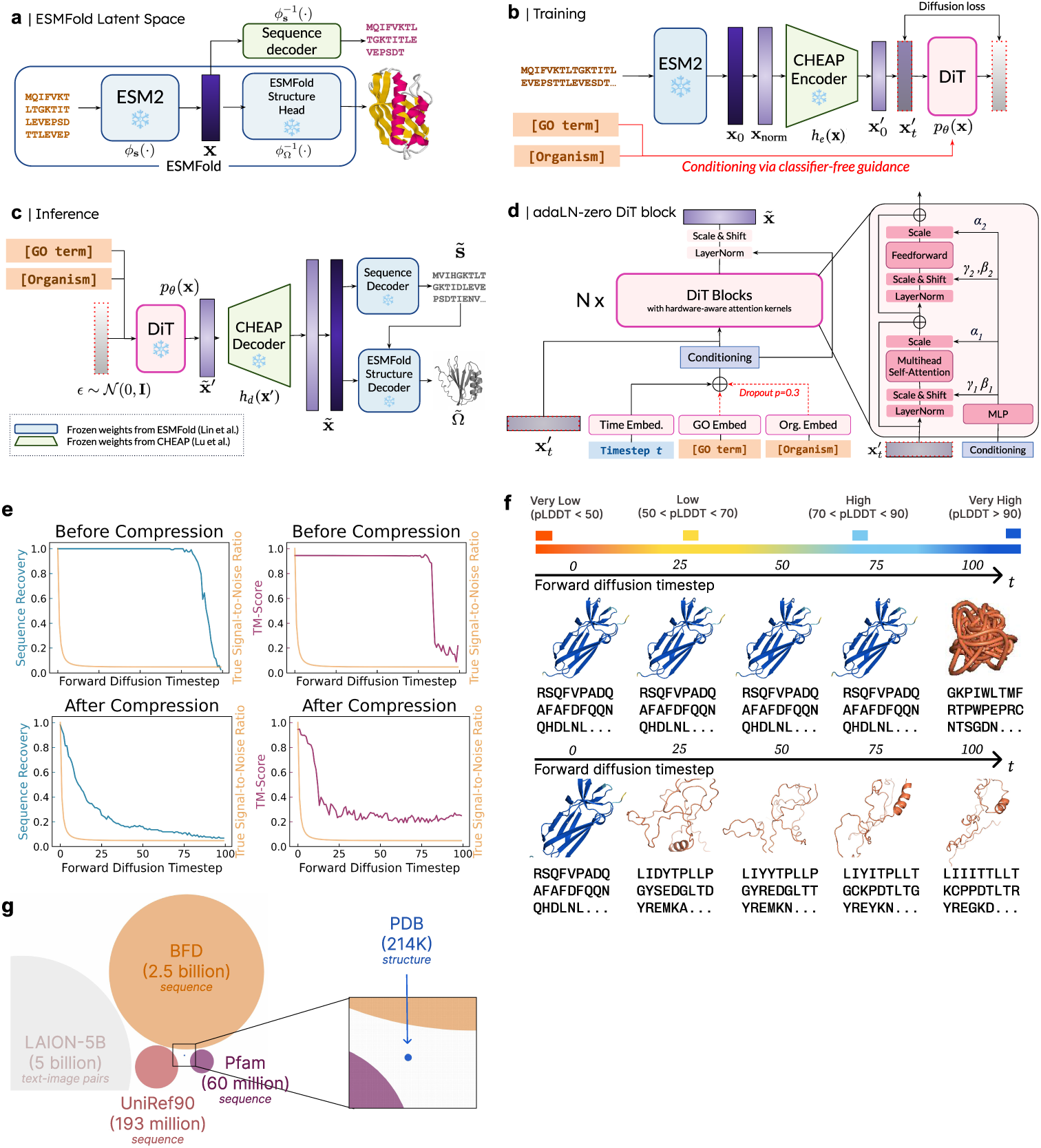
(a) Defining a joint latent embedding of sequence and structure. The intermediate outputs of highly capable protein folding models contain all information for structure, but require only sequence to obtain. This schematic of ESMFold [9] illustrates the latent embedding **x** used to define a joint representation of sequence and structure. We use an additional sequence decoder from [25]. The frozen sequence and structure decoders allow us to map this joint embedding back to sequence and structure. We consider this a joint embedding of structure and sequence; learning to generate this embedding thus allows for simultaneous multimodal generation. **(b) Training.** During training, we use denoising diffusion probabilistic models (DDPM) [30] to generate the latent embedding. To improve model performance and mitigate “unsmoothness” in the latent space, as further described in Figure 1e and f, we first compress the embedding using the compression encoder in the autoencoder introduced in CHEAP [25] from 1024 to 32 channel dimensions, with a 2× shortening factor. **(c) Inference.** During inference, we generate a latent embedding from the learned diffusion model, then use the CHEAP uncompression decoder. To obtain the sequence, we use a frozen sequence decoder from CHEAP (see Figure 1a). To obtain structure, we reattach the frozen ESMFold structure decoder head, using the amino acid identities provided by the sequence decoder, to obtain the all-atom structural generation. **(d) Architecture and conditioning.** PLAID uses the Diffusion Transformer (DiT) architecture with adaLN-zero [27] blocks to incorporate conditioning information. Classifier-free guidance is used to incorporate both the function (i.e., GO term) and organism class label embeddings. DiT block schematic adapted from Peebles and Xie [27]. **(e) Latent space regularization (quantitative).** Diffusion models rely on learning to undo noise added according to a noise schedule. We find that the perturbations here do not map to noise in the structure and sequence spaces after applying the decoder. As a result, directly learning a diffusion model in this space produced poor initial results. The orange line denotes true noise schedule signal-to-noise ratio. Sequence recovery refers to per-token identity between the original sequence and the decoded sequence (after latent space noising). TM-score measures structure similarity between the unnoised structure and the decoded structure after latent space corruption. After compression and normalizing the latent space, to fix the massive activations reported in Lu et al. [25], the resulting corruptions in sequence and structure space more closely match the true signal-to-noise ratio added to the latent space. **(f) Latent space regularization (qualitative).** Examining results reported in Figure 1e visually. Without compression, noise in the latent space does not result in corrupted structures, as visualized by the high pLDDT [60]. **(g) Drawn to scale comparison of database sizes**. Compared to structure databases such as the PDB, sequence databases can be 2-4 orders of magnitude larger, and furthermore have better coverage of unstructured protein space which can also be useful to be represented when generating proteins (e.g. intrinsically disordered regions). For reference, we also show the size of Internet-scale datasets [63], to show the difference in data availability for the PDB compared to the text-to-image setting, from which diffusion model techniques originated.

### 2.1 Capturing multimodal information in protein folding models

A central motivation for PLAID is that the internal representations of large folding models inherently jointly represent sequence and structure, since it is obtainable from sequence, but contains all the information necessary for highly accurate structure prediction. Therefore, we learn to generate embeddings directly from this latent space. Following Lu et al. [25], we extract such an embedding from ESMFold [9], immediately upstream of its structure module (Figure 1a).

This affords several advantages. Directly learning to generate a joint representation of structure and sequence mitigates the need for multi-stage processes common in all-atom generation [18, 19], whereby an inverse folding stage is necessary after backbone generation. Further, structural databases are small and biased toward crystallizable proteins, whereas sequence repositories are orders of magnitude larger (Figure 1g). Since generative models inherently interpolate from their the distribution defined by their training data, this allows the model to sample from a more natural evolutionary-scale database. Additionally, sequence databases are richly annotated, allowing for new axes of generative control (Section 2.3). By sampling from the latent space of a pretrained protein folding foundation model, PLAID exploits the structural priors encoded in the weights of pre-existing and highly capable models. This paradigm can be broadly applied to any predictive model mapping a more abundant data source to another, whereby learning the intermediate latent space enables generating the less abundant data source while being able to instead interpolate from the expanded coverage enabled by the more abundant data source.

### 2.2 Latent compression for stable diffusion

To learn a generative model over the latent space, we turn to the highly successful diffusion paradigm [30, 31] (Figures 1b and c, Section 6.3). However, we find that learning a generative model directly on the original latent space produces poor results and unstable learning (Extended Data Figure 3). To diagnose this, we observed that the signal-to-noise ratio (SNR) at a given diffusion timestep does not map smoothly back to sequence or structure space (Figure 1e, f). Even when an aggressive noise schedule is used, the sequence and structure remain uncorrupted until the final forward diffusion timesteps, meaning most sampled timesteps are trivial for learning.

Lu et al. [25] report that protein folding model latent spaces exhibit “massive activations” whereby some channels have magnitudes 3000× larger than the median channel value, consistent with those observed in Transformer models broadly [32]. We hypothesize that this is causing the non-smooth mapping from the latent to the structure space. We therefore use the compression autoencoders in Lu et al. [25] that normalizes and smooths the latent space. An intuition for this approach comes from high-resolution image synthesis literature, where it is often necessary to first compress images before learning a diffusion model; this further has the advantage of accelerating sampling speeds (Extended Data Table 4.)

We reduce the embedding dimensionality (1024→32 channels) and downsamples length by 2×, and find that learning this compressed latent space drastically improves performance. After compression, the signal-to-noise ratio (SNR) is more similar between the true noise schedule when added in latent space and when mapped back to the sequence and structure space (Figure 1e, f). At inference time, generated latent embeddings are first decoded with the autoencoder before passing to the sequence and structure decoders (Figure 1b).

### 2.3 Compositional control by function and organism

Many deployed video and image generation models are controlled via natural language interfaces, affording greater controllability by the user. Analogously, direct specification of desired biochemical properties through language-based prompts can afford similar controllability in protein engineering. Because PLAID is trained on sequence-scale databases rather than structural corpora, it benefits from the dense functional and taxonomic annotations available, providing a path towards this goal. Consequently, we train PLAID with classifier-free guidance using 2,219 Gene Ontology (GO) terms [26] and 3,617 unique organisms. Both can be compositionally specified, with separate guidance weighting.

Unlike prior approaches that achieve control through explicit structural motif scaffolding, PLAID can generalize beyond motif-based constraints, or specifying active sites *a priori*. This distinction is particularly important for enzyme design and other functions where non-adjacent residues coordinate interactions. In addition, PLAID introduces organismal conditioning, which reflects a key dimension often neglected in generative modeling. In practical drug discovery, successful design depends not only on binding, but also on developability. Conditioning on organismal context acknowledges immunogenicity constraints for antibody design, sequence- and structure-level differences between prokaryotic and eukaryotic proteins serving similar functions, and that expressibility might be specific to certain cell lines.

PLAID is implemented as a fully Transformer-based Diffusion (DiT) [27], enabling efficient scaling beyond two billion parameters through optimized attention kernels [29]. Combined with massive augmentation from training on sequence-scale data, PLAID makes use of modern observations in machine learning that massive capability gains often arise from scaling parameters and data.

## 3 In silico results

### 3.1 Unconditional generations

To assess performance as a function of chain length, we generated 64 samples per target length across 64, 72, 80*, . . . ,* 512, yielding *n*=3,648 proteins, following prior length-stratified evaluations [15, 18, 20]. Details of designability metrics (i.e. cross-consistency and self-consistency), diversity, and novelty metrics are defined in Supplemental Information Figure 1 and Extended Data Table 6. Figure 2 shows length-stratified designability and biophysical plausibility, and Figure 3 shows diversity results.

**Fig. 2:**
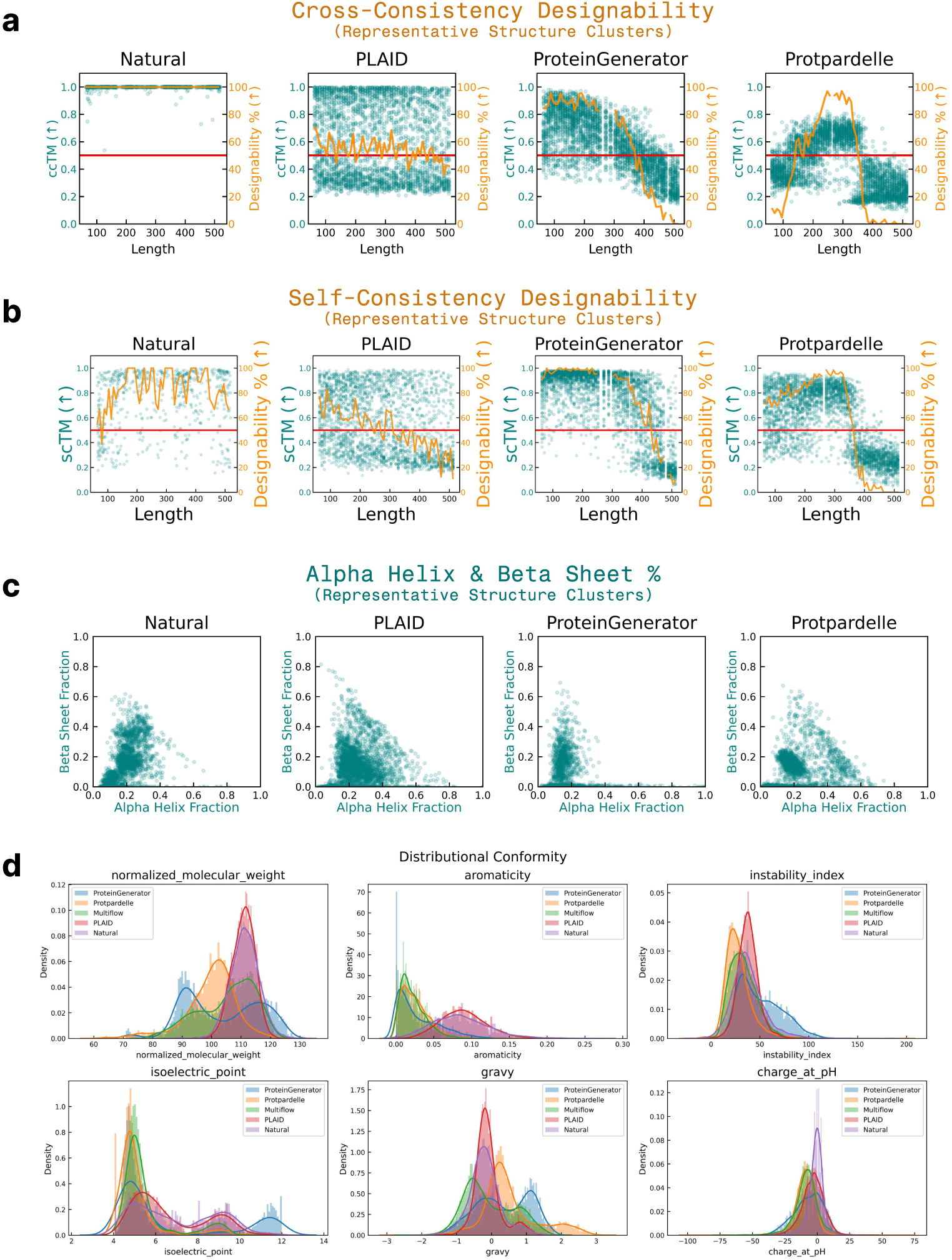
*In silico* comparison of generation quality. Tabular version of these results are shown in Extended Data Table 1. Analysis performed on 64 samples per length interval. The red line indicates the TMScore *>* 0.5 threshold for designability. **(a)** Designability defined by cross-consistency, which assesses if co-generated sequence and structure accord with each other. Teal points (left axis) denote ccTM values samples in the given length interval. Samples are first clustered via foldseek easy-cluster, to account for the quality-diversity trade-off commonly observed for generative models. Orange line (right axis) measure the fraction of samples at that length interval which are designable (defined as ccTM *>* 0.5). In the unconditional generation setting, PLAID is able to generate more designable sequences at longer sequence lengths compared to baselines. **(b)** For completeness, we also report the designability defined by self-consistency, as is common in unimodal structure generation works. See Extended Data Table 6 and Supplementary Information Figure 1 for more information on how metrics are defined. **(c)** Measuring fractions of alpha helix versus beta sheet proportions in generations after clustering (to consider diversity). Traditional methods often struggle to generate beta-sheets during unconditional generation; this can obfuscate designability metrics, since alpha-helical regions generally have higher consistency. In contrast, PLAID unconditional generations produce more diverse samples with good representation of beta-sheet regions. **(d)** *In silico* evaluation of quality by examining biophysical similarity to natural proteins, which is shown to accord well with wet-lab succcess [35]. PLAID generations accord more closely with natural proteins biophysically. Properties are detailed in Supplemental Information B.6. Charge at pH is measured for pH=7.

**Fig. 3:**
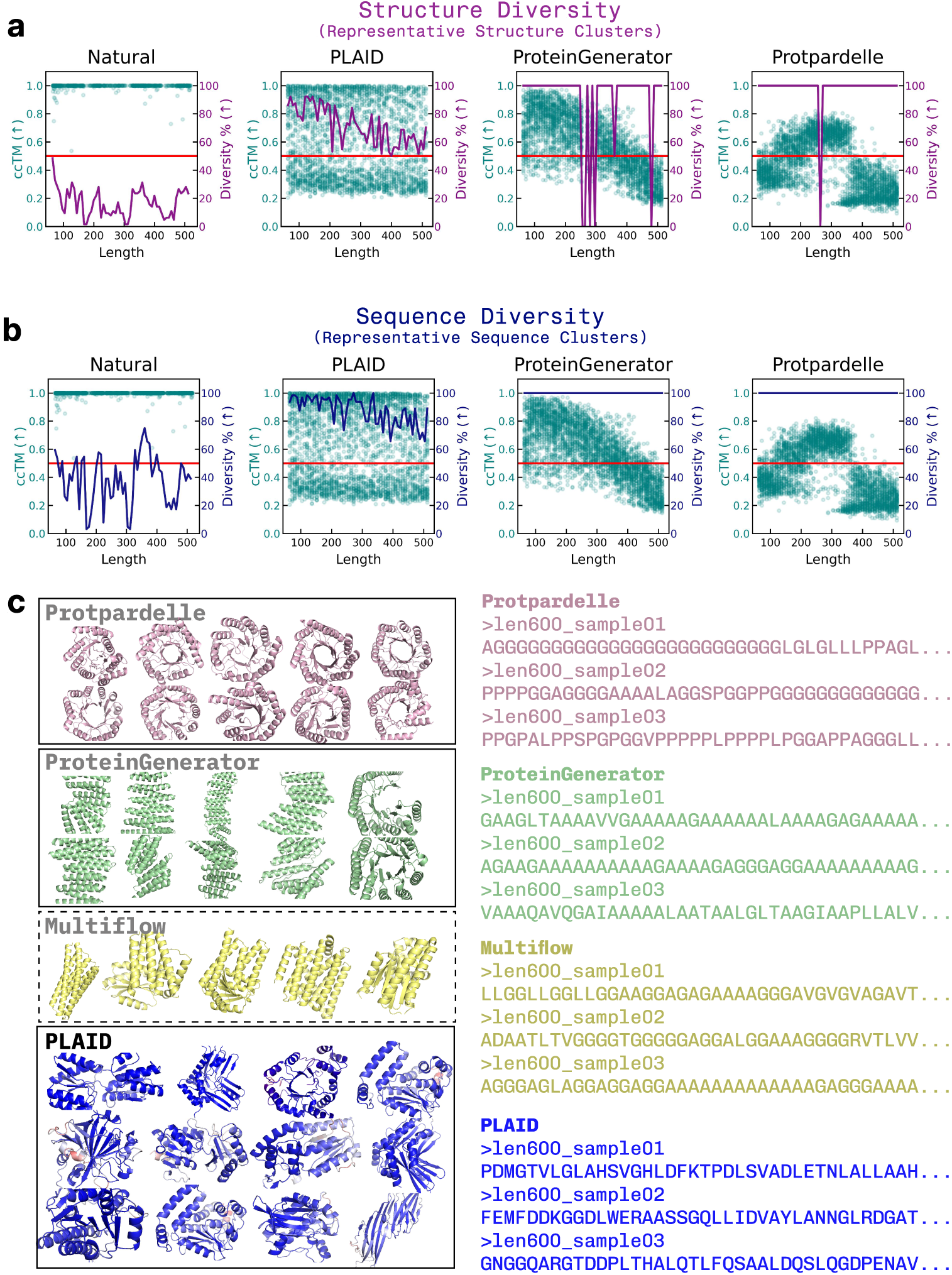
Examining diversity for unconditional generations. Similar to Figure 2, we stratify by length with 64 samples per interval between 64 to 512 residues, where each teal point is a single unique example after clustering. **(a)** Structure diversity percentage at each length is defined by the number of unique structure clusters (using foldseek easy-cluster using default settings), plotted in purple (right axis). For reference, the cross-consistency TM-Score for generations are also plotted. For baselines, at certain lengths, mode collapse is observed, where generations suffer from severe lack of diveresity. **(b)** Sequence diversity percentage is the number of unique sequences calculated using mmseqs easy-cluster. Similar to Figure 3a, ccTM is plotted for reference. **(c) Qualitatively examining mode collapse in baselines.** To shed light on quantitative metrics, we qualitatively examine generations. For structure generations (visualized for length of 256), as shown in Figure 3, samples often suffer from mode collapse. For sequences (visualized for length 600), baseline samples often suffer from repeats, which is not observed for PLAID samples.

#### PLAID better balances quality and diversity at longer sequence lengths

As shown in Figure 2a and b (cross-consistency and self-consistency, respectively), the decline in structural quality with increasing length is markedly less pronounced for PLAID than for baselines. Secondary-structure composition further highlights these differences: PLAID’s distributions track natural proteins more closely, whereas existing models frequently underproduce *β*-sheet rich folds (Figure 2c).

Aggregated over lengths, cross-modal agreement metrics (Extended Data Table 1) are highest for PLAID, consistent with the fact that PLAID directly samples from the joint distribution of sequence and structure, as compared to baselines which sample either structure or sequence, and then predict the other modality. We quantify the quality–diversity trade-off by counting distinct sequence and structure clusters that are *designable*, defined as ccRMSD *<* 2°A. Among all-atom models, PLAID yields the largest number of distinct, designable clusters in both spaces (Extended Data Table 2). While some prior methods appear more “novel” by nearest-neighbor criteria, these scores can be inflated by low-quality outputs that are merely dissimilar to natural proteins.

#### PLAID samples have high distributional conformity to natural proteins

As shown in Figure 2d, across molecular weight, aromatic amino-acid fraction, dipeptide-based instability index [33], hydropathy (GRAVY) [34], isoelectric point, and net charge at pH 7, PLAID better recapitulates the distributions observed in natural proteins than baselines. Prior work connects such conformity with improved expressibility and practical success [35]; detailed Wasserstein analyses are provided in Appendix B.6.

#### PLAID avoids length-specific mode-collapse

Figure 3a examines structural diversity, overlayed with quality. For baselines, at specific lengths, mode collapse is observed, whereby outputs are concentrated into a small set of structures. Figure 3b similarly examines this for sequence diversity. In Figure 3c, qualitative demonstration of mode collapse (e.g. TIM-barrels for Protpardelle [18] at length 256) is shown. Additionally, we see that for longer sequences, baselines are prone to generate sequence region repeats, a qualitative artifact that might go uncaptured by perplexity and sequence recovery metrics (Extended Data Figure 1b).

### 3.2 Conditional generations

#### PLAID recapitulates active-site side chains

Across a wide panel of enzymatic functions, PLAID-generated structures recover defining active-site geometries, including metal coordination spheres, heme-pocket scaffolds, and placement of catalytic acid/base residues (Figure 4a). For each design, we identify the closest ligand-bound structure by Foldseek [36] and examine active site correspondence. Although global sequence identity to the nearest experimentally solved structure is typically less than 40%, active site side-chain residue identities and orientations closely match those observed in crystallographic complexes. This suggests that the model has learned information about residues involved in active sites associated with the function, without directly memorizing the sequence. Importantly, this ability to generate enzymes does not involve knowing the indices *a priori*, and many active site residues that are correctly recapitulated are not directly adjacent to each other. Beyond enzymes and membrane proteins, prompts corresponding to transferases, ligases, hydrolases, and receptor-binding proteins produce folds appropriate for the functions (Figure 4c).

**Fig. 4:**
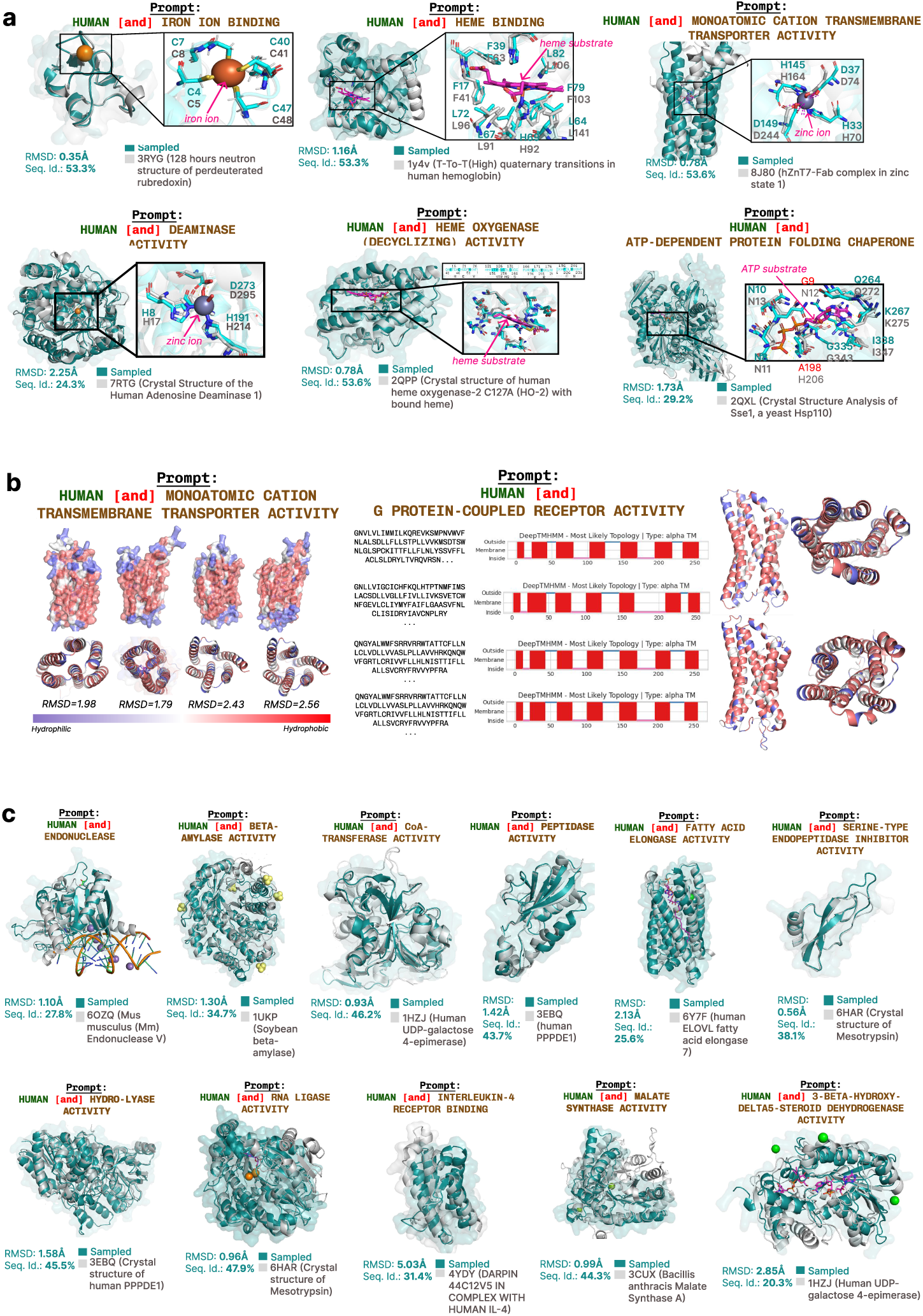
Function-conditioned generation vignettes. (a) Examining active sites. For each Gene Ontology keyword prompt, PLAID samples a *de novo* sequence–structure pair. Using Foldseek [36], a structure-based search is done to find the closest PDB sample that was resolved in-complex with a ligand, and super-imposed. Function-conditioned generations recover defining active-site geometries across diverse enzyme classes despite low global sequence identity. Active sites exhibit correct placement and orientation of catalytic residues, metal-coordination geometries, or substrate-binding pockets, even when sequence similarity is *<* 40%. Notably, active site residues may not be directly adjacent, and the model does not rely on explicit motif scaffolds or structure templates. **(b) Membrane proteins.** PLAID generates membrane proteins with physically realistic hydropathy organization and family-specific topologies directly from functional prompts. Conditioned on transmembrane-transporter or GPCR activity, the model produces multi-helix topologies with appropriate segregation of hydrophobic and polar residues, and seven-helix GPCR-like architectures, despite receiving no 3D structural supervision during training. This suggestions that PLAID has absorbed biophysical priors about amino acid hydrophobicity and implications for membrane-embedded proteins as a co-generation model of sequence and structure. **(c) Additional conditional generations.** Additional examples across enzymatic, binding, and regulatory functions highlight compositional control. Prompts specifying enzymatic activity (for example, ligase, hydrolase, transferase), molecular function (for example, inhibitor activity, interleukin-4 receptor binding), or metabolic roles produce structures that match the expected fold class despite having low sequence identity.

#### Transmembrane protein generations have expected hydrophobicity patterns

When conditioned on transmembrane transporter activity, PLAID generates multi-pass helical bundles with hydrophobic exterior surfaces and polar channel interiors (Figure 4b). Generations prompted by G protein–coupled receptor (GPCR) activity yield seven-helix topologies with the expected helix packing and amphipathic patterning. Applying DeepTMHMM [37] to generated GPCR-prompted sequences, it is similarly able to predict a seven-helix topology, as anticipated. These behaviors emerge despite lacking explicit structural supervision during diffusion model training.

Together, these analyses suggest the possibility of a new interface for controlling function beyond explicit motif scaffolding. However, for completeness, we demonstrate how PLAID can be used for motif scaffolding in Extended Data Figure 1a.

#### Organism-prompted generations cluster similarly in t-SNE space

Embedding visualizations of organism-conditioned generations (Figure 5) show separation concordant with phylogenetic distance (e.g., *Glycine max* vs. *E. coli* ) and tighter proximity for closely related taxa (human vs. mouse). For human-conditioned designs in Figure 4, 66.5% have a nearest mmseqs easy-search neighbor that come from *Homo sapiens*. For reference, we also repeat the same analysis for real proteins from the same organisms, randomly sampled from UniRef50. These organism-level distinctions are mostly sequence-driven, and are therefore uniquely unlocked by PLAID’s co-generation capabilities.

**Fig. 5:**
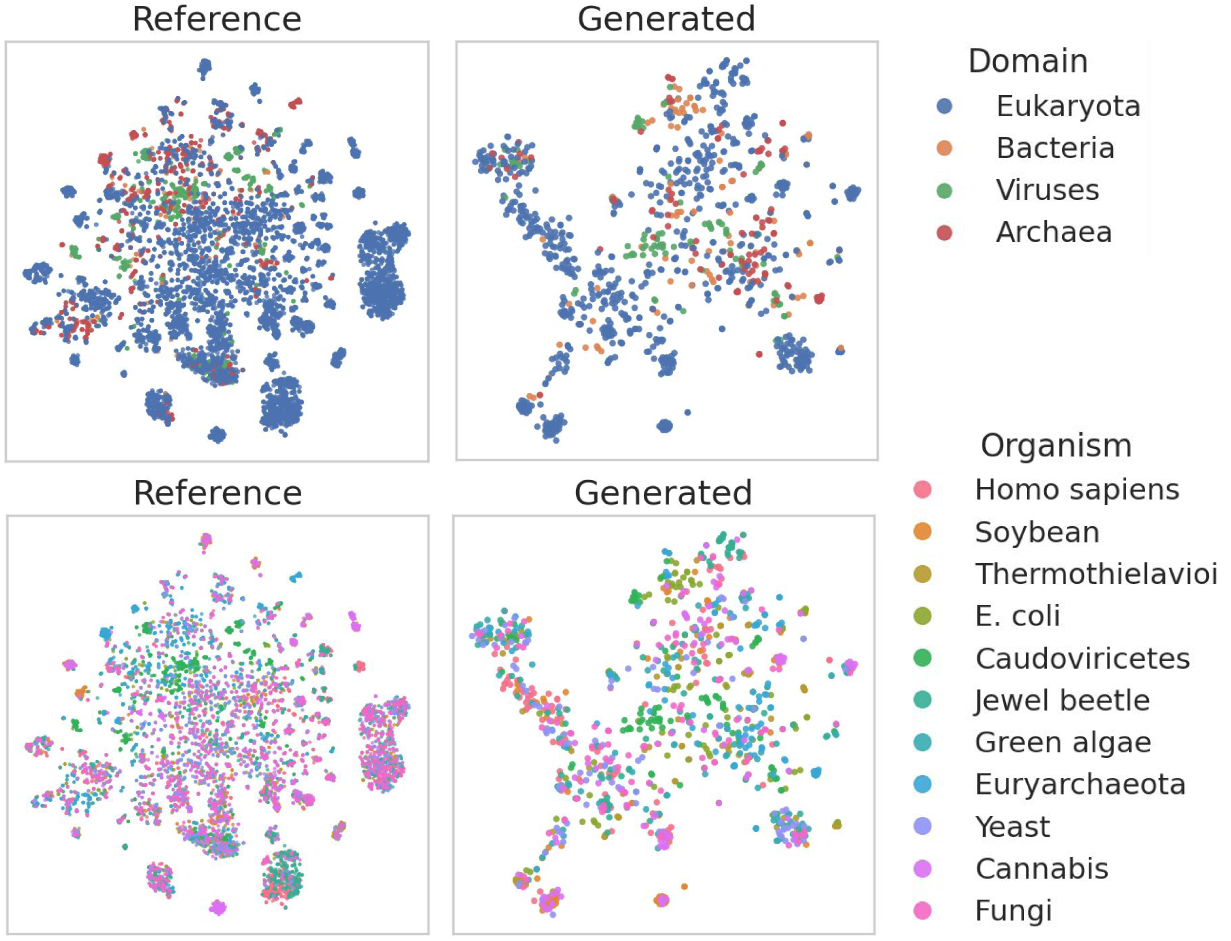
Taxonomic conditioning yields organism-specific structure–sequence distributions. t-SNE projections of latent embeddings for natural UniRef50 proteins (left) and PLAID-generated proteins conditioned on taxonomic prompts (right). When prompted with domain-level (top) or organism-level (bottom) labels, PLAID produces sequence–structure pairs that occupy separable regions of latent space mirroring the phylogenetic organization of natural proteins. Because taxonomic differences mostly manifest at the sequence-level rather than in structure, this result demonstrates that PLAID’s joint sequence–structure generation internalizes organism-specific sequence signatures in addition to its atomic-level precision in maintaining function associations.

## 4 Experimental validation of heme-binding proteins

To experimentally verify the functionality of the designs, we generated heme binding proteins and tested them for both expression and heme binding. Heme binding proteins play a crucial role in a wide range of biological processes essential for life such as oxygen transport, electron transfer, and a plethora of catalytic reactions essential for life such as signaling pathway, detoxification pathways and normal metabolism [38–40]. Moreover, heme enzymes have been engineered to perform non-native chemical transformations with synthetic utility [41]. They have therefore often been targets for *de novo* design [42–46].

### 4.1 Sample Generation

Heme-binding proteins were generated by prompting using the GO keyword “heme-binding” and the target organism *E. coli*, chosen as the expression host. For the generation, two lengths (120 and 160 amino acids) were selected because of an higher overall designability of those generations. The corresponding PLAID-generated proteins are referred to as“Globin” and “H-NOX” folds respectively, which was determined by using Foldseek to find protein folds with matching structures in the PDB, showing that the PLAID-generated protein structures of length 120 largely correspond to the fold of proteins from the Globin family, and those of length 160 correspond to the fold of proteins from the H-NOX family. To further evaluate PLAID’s backbone generation, the 120-residue PLAID-generated structures that were deemed designable were also inverse-folded using the standard ProteinMPNN, and these are referred to as “IF Globin” proteins.

After generating 1000 sequences for each category (Globin, H-NOX, IF Globin), we filtered them down to retain only those considered designable and containing an axial histidine in the correct position for heme coordination (Section 6.8 and Supplemental Information Figure 1, Figure 6a). These proteins showed varying levels of quality and novelty, as measured by structural folding confidence and sequence identity to known proteins (Figure 6b–d). We then selected proteins with high pLDDT and ipTM values from diverse clusters (Section 6.8 and Supplemental Information Figure 1), resulting in 17 IF Globin, 34 Globin, and 9 H-NOX proteins for experimental validation.

**Fig. 6:**
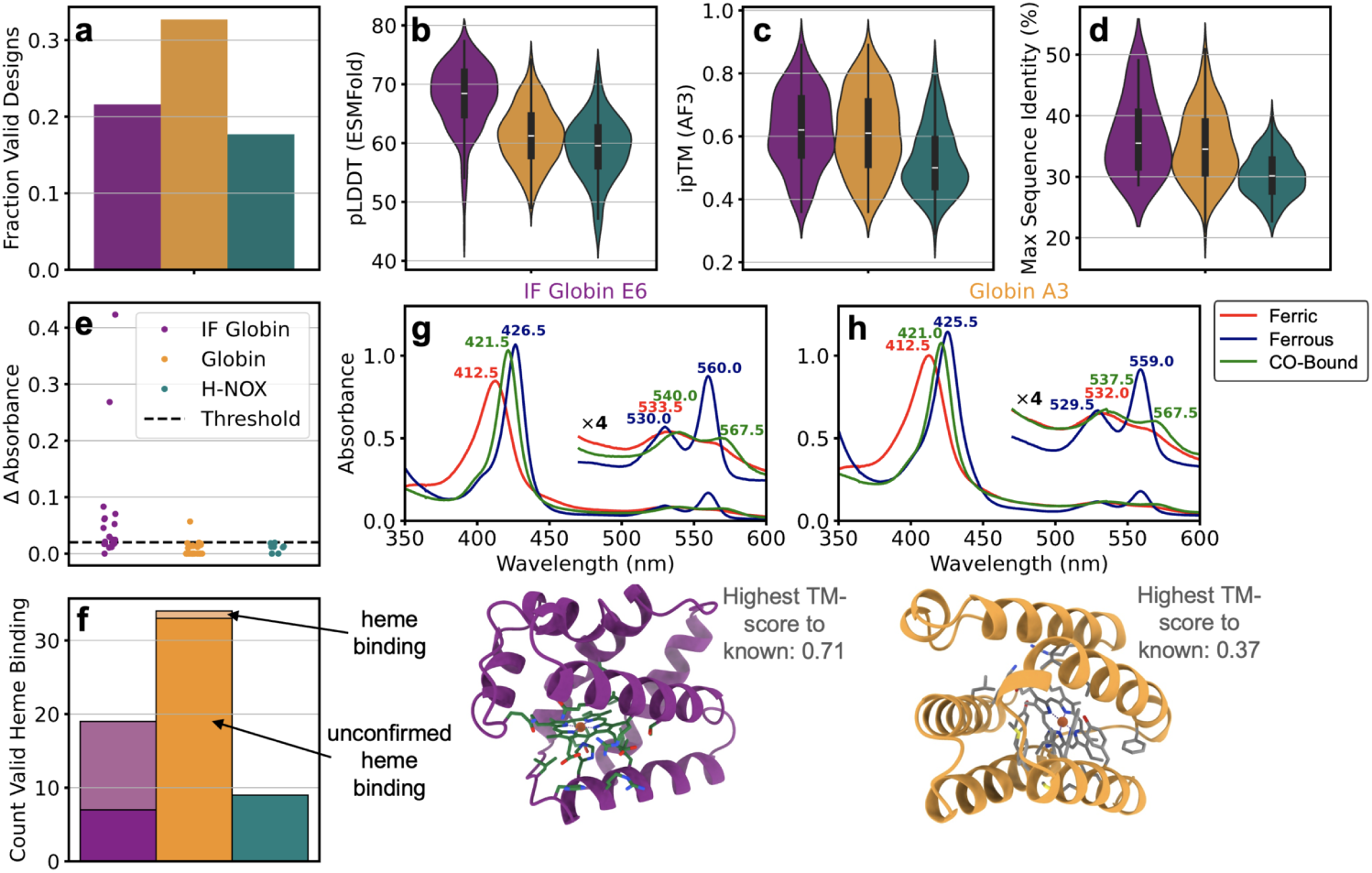
Experimentally validated heme binding proteins through function-conditioned generation from PLAID. **(a)** Fraction of all PLAID-generated sequences that passed initial computational filters for designability and correct heme axial ligand: for inverse-folded (IF) globin sequences of length 120, globin sequences of length 120, and H-NOX sequences of length 160. Computational metrics of sequences that passed the initial computational filters: **(B)** pLDDT of the protein folded without heme from ESMFold, **(C)** the ipTM confidence of heme relative to protein as folded by AlphaFold3 (AF3), and **(D)** the novelty of sequences as measured by the maximum sequence identity to a know protein in Uniref90, from Diamond BLAST. Based on ESMFold, nearly all generated sequences adopted a known protein fold of a Globin or H-NOX protein, respectively. **(E-F)** Among the proteins generated by PLAID and selected for experimental validation, many of the inverse-folded globins showed heme binding, and a few of the directly generated sequences from PLAID showed heme binding, as measured by the change in absorbance at 428 nm between ferric and ferrous forms, from wet-lab experiments using lysate. Absorbance spectra for **(G)** an inverse-folded globin protein (E6) and **(H)** a globin protein (A3) generated from PLAID, showing a change in absorbance under purified protein wet-lab conditions, which confirm heme binding. Respective AF3-predicted structures are shown, with the Globin sequence in A3 (generated directly from PLAID) showing high overall structural novelty (TM-score of 0.37) compared to the closest known protein in the PDB.

### 4.2 Experimental characterization

Following expression of the generated proteins in *E. coli*, their absorbance spectra were measured in cell lysate with and without reduction. This revealed that a significant fraction of the IF Globin proteins successfully bind heme, while only some Globin-type proteins did so. None of the H-NOX proteins showed detectable heme binding. The absence of signal may result from lack of inherent binding capability or overall expression level. To further validate these findings, one representative protein from each promising group were selected for purification: IF Globin E6 and Globin A3. After purification, Globin A3 yielded 2.5 mg/L of culture, whereas IF Globin E6 yielded 122.5 mg/L of culture. Furthermore, Globin A3 exhibited some precipitation upon concentration, suggesting lower stability compared to IF Globin E6. In contrast, both proteins displayed clear heme-binding spectral signatures (Figure 6g-h) consistent with previously described bis(histidine) heme bound proteins [43, 47]. Together, these results validate that PLAID can be functionally-prompted to generate heme-binding proteins.

## 5 Discussion

We propose PLAID, a model for controllable generation of all-atom proteins, which can be prompted by functional keywords and taxonomy. By training in the latent space of a protein folding model, we can obtain sequence-scale coverage while generating with atomic-level precision that recapitulate enzymatic active sites. Experimental validation shows that heme binders can be generated with low sequence similarity to known proteins.

Although we use the latent space of ESMFold [9] in this work, the method can be applied to any prediction model from a less abundant to a more abundant modality to expand data coverage. It is forward-compatible with increasingly multimodal prediction models [7, 46, 48–50], and diffusing in the latent space and using frozen decoders from these models can allow for generating more modalities than all-atom structure. The ability to train on this more abundant data source (and therefore have access to more annotations) open the door for improved control when prompting this generative model. For example, while taxonomy is not a direct developability metric, controlling for organismal origin is a step toward capturing contextual constraints (e.g. host-dependent expression environments) that are relevant in applied protein engineering.

While latent generation is beginning to be explored for protein generation, PLAID uniquely explores how it can be used to augment usable data, and explores compression mechanisms. Compression allows for faster generation, and opportunities to pre-filter generated latent embeddings prior to decoding. We hypothesize that the compression mechanism and therefore being able to train on longer sequences contributed to our improved performances at longer protein sequence lengths.

By expressing the designed proteins in *E. coli*, we showed that some PLAID sequence-structure co-designed proteins exhibit spectroscopic signature characteristic of heme-bound proteins, with generated folds similar to proteins adopting Globin or H-NOX folds. Noteworthy, while the Globin-like folds were functional, the H-NOX–like designs were largely inactive, highlighting that different folds or different sequence lengths may not respond similarly to the same generative conditioning. Interestingly, inverse folded sequences demonstrated better overall expression and stability, corroborating findings elsewhere in the field Sumida et al. [51]. Generated heme sequences have low sequence identity relative to known proteins, indicating genuine novelty rather than mere memorization. All of the directly generated structures from PLAID adopt the same overall fold as known Globin and H-NOX proteins. Together, these results demonstrate that PLAID can conditionally generate functional, novel, and diverse proteins guided by GO terms.

A limitation of PLAID is that performance can be bottlenecked by the prediction model from which the frozen decoders are derived. With explicit fine-tuning for latent generation (e.g., training CHEAP and the structure decoder end-to-end), model performance might be improved. Furthermore, since the current structure decoder is deterministic, it does not sample alternative conformations. A solution is to use a decoder that returns a distribution over structural conformations instead. Additionally, the GO term one-hot encoding used here does not take into account the hierarchical nature of the Gene Ontology vocabulary. This can be fixed by using a multi-class conditioning scheme instead. Finally, the classifier-free guidance scale can be separated for the organism and function conditions, since the two may require different guidance strengths in real-world use cases. These limitations will be examined in future work.

## 6 Methods

### 6.1 Notation

A protein is composed of amino acids. A protein sequence 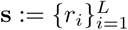 with length *L* is often represented as a string of characters, with each character denoting the identity of an amino acid residue *r_i_* ∈ ℛ, where |ℛ| = 20. Each unique residue *r* can be mapped to a set of atoms 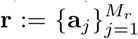, where each **a***_j_* ∈ ¡^3^ is the 3D coordinate of an atom, and the number of atoms *M_r_* may differ depending on the residue identity. A protein structure 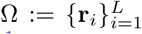 consists of all atoms in the protein and therefore implicitly contains **s**.^1^ To reduce complexity, protein designers sometimes work only with backbone atoms Ω_backbone_ ⊂ Ω only, which consists of [*C_α_, C, N*] atom repeats. These are sufficient to define the general fold.

### 6.2 Defining a Joint Distribution of Sequence and Structure

Our goal is to characterize a joint embedding of structure and sequence information *p*(**x**) over X, such that there exist mappings **x** = *ϕ***_s_**(**s**) and **x** = *ϕ*_Ω_(Ω). To do so, we use the latent space of protein folding models, allowing us to repurpose information from these *predictive* models for *generation*. The trunk of the model provides **x** = *ϕ*_ESM_(**s**), and the structure module head provides Ω = *ϕ*_SM_(**x**). If we consider an implicit inverse function of the structure module such that 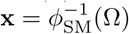, then we have **x** = *ϕ***_s_**(**s**) = *ϕ*_Ω_(Ω), and **s** and Ω can be mapped to the same **x**.

#### Overview of ESMFold

Briefly, ESMFold [9] has two components: a protein language model component **x** = *ϕ*_ESM_(**s**), and a structure module component Ω = *ϕ*_SM_(**x**) that decodes these latent embeddings into a 3D structure. For **x** ∈ ℝ*^L^*^×1024^, we use the layer outputs representation just prior to the structure module (pseudocode provided in Appendix B.1). This is because when ESMFold is used at inference time, the pairwise input is initialized as zeros, such that the sequence input contains all information necessary for structure prediction (Figure 1A).

#### Latent Generation

Our goal is to learn *p_θ_*(**x**) ≈ *p*(**x**), where *θ* is the set of parameters of the model learned through diffusion training (Figure 1C). Then, after training, we can sample **x̃** ∼ *p_θ_*(**x**) (Figure 1C). To do so, we use diffusion models [30, 31] with some modifications (described in ablation Table 3).

### 6.3 Diffusion Model Training

Ablations for the training details are in Table 3. We adopt the discrete-time variance-preserving diffusion formulation [30]. A “clean” latent embedding **x**_0_ ∼ *p*(**s**, Ω) from above is gradually corrupted by adding Gaussian noise over *T* timesteps. A key result from Ho et al. [30] is that during this forward diffusion process

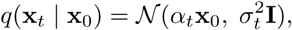

we can express the noised latent embedding **x***_t_* in closed form given the uncorrupted **x**_0_ as

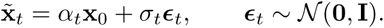

#### Sigmoid noise schedule

Prior work [53] show that using a sigmoid noise schedule rather than the cosine [54] or linear [31] schedules can improve performance by shifting weight away from the tails of the schedule. We similarly use the Sigmoid schedule

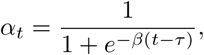

where *β* and *τ* control the shape of the schedule.

#### Min-SNR loss reweighting

Diffusion model training sometimes suffers from slow convergence due to the conflicting optimization directions between timesteps. For the signal-to-noise (SNR) ratio 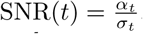, a reweighting strategy is applied to each timestep. This effectively treats each step as a separate “task”, and the diffusion training process overall as a multitask problem, whereby harder tasks are weighted more heavily in the loss function. We adopt the Min-SNR-*γ* formulation introduced in [55], such that at each timestep, the SNR is clamped to a maximum value *γ >* 0 as

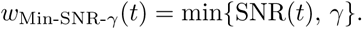

#### Velocity prediction objective

Salimans and Ho [56] propose to replace the *L*_simple_ objective with a velocity-based objective. Rather than minimizing

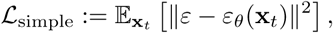

we minimize

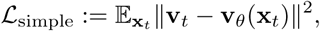

where 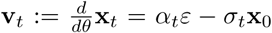. This is done to improve training stability and sample quality, as it allows for a more direct optimization of the denoising process. We hereafter refer to the diffusion model as *v_θ_*(·).

#### Self-conditioning

We find that using self-conditioning [57**?**] both improves *v_t_*-prediction performance and sample quality. In self-conditioning, the denoiser also takes its own previous prediction as input as *f_θ_*(**x***_t_,* **x̂**_0_*, t*), allowing the model to iteratively refine its output. Within *v_θ_*(·), we have a linear projection layer that combines 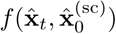 to have the same dimensions as **x̂***_t_*.

Together with Min-SNR-*γ* weighting, *v*-prediction, and **x**_0_ ∼ *p*(**s**, Ω) defined in Section 3.1, the training objective becomes:

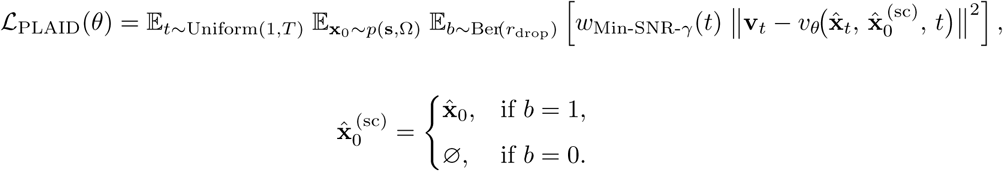

### 6.4 Latent Space Compression

In our initial experiments, we found empirically that directly learning *p_θ_*(**x**) without compression performed poorly (Figure 5). We suspected that this might be due to the high dimensionality of **x** ∈ ℝ*^L^*^×1024^. In comparison, Rombach et al. [58] use a latent space of size **z** ∈ ℝ^32×32×4^. We were therefore inspired to mirror works in high-resolution image synthesis and compress the latent space before learning the diffusion model.

Compression is performed using the CHEAP autoencoder [25], *h_e_*(**x**). After examining the bit–distortion rate results in Lu et al. [25], we choose to compress from 1024 → 32 channels and downsample by 2× along the length. For all details in the training section, all instances of **x**_0_ are replaced with **x**^′^ = *h_e_*(**x**).

### 6.5 Sampling Latents and Decoding Sequence and Structure

#### Latent Sampling

All results in the main text use the DDIM sampler with 500 timesteps unless otherwise noted. During sampling, we derive **x̂***_t_* from **v̂***_t_* as

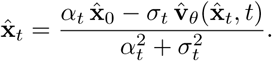

We also apply an exponential moving average (EMA) to model parameter weights, as this has been shown to improve sample quality [30].

#### Decoding Sequence and Structure

An implicit inverse mapping of ESM2 to get 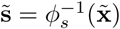 can be trained with a validation accuracy of 99.7% on a held-out partition of UniRef [9]. To obtain sequence and structure, we apply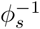 and the frozen ESMFold decoder.

### 6.6 Compositional Conditioning by Function and Organism

In classifier-free guidance (CFG) [59], a single network operates both conditionally and unconditionally by randomly replacing the class *y* with the null token ∅ during training with some probability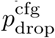. We sample 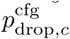 and 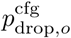 independently.

At inference, for each *x_t_* we compute

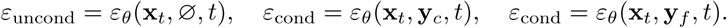

With guidance scale *w* ≥ 1 (defaults to *w* = 3 in our experiments), the guided prediction is:

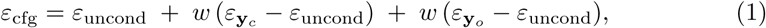

### 6.7 Architecture

We use the Diffusion Transformer (DiT) [27]. In early experiments, we found that allocating available memory to a larger DiT model was more beneficial than using triangular self-attention [60]. We train two variants of the model: a 2B version and a 100M version, both with the memory-efficient attention implementation in xFormers, using float32 precision. Due to the way that xFormers is implemented, at inference time, sequence lengths must be a multiple of 4. All other models are trained with mixed precision (bfloat16 and float32). A learning rate of 1e−4 was used, with cosine annealing applied over 800,000 steps. The xFormers memory-efficient attention kernel requires input lengths to be a multiple of 4. Since we also apply an upsampling factor of 2, the actual inference length must be a multiple of 4. During training, the maximum sequence length we use is 512, based on the distribution of sequences in Pfam and a shortening factor of 2 based on results in Lu et al. [25].

### 6.8 Conditional Generations

#### In silico generation

Evaluation of conditional generations introduces a conundrum where structure and sequence should be highly conserved when they exhibit similar functions, and thus similarity to known proteins should indicate successful conditioning. However, in machine learning literature, works often prefer generations being *different* from known proteins to emphasize novelty or lack of training data memorization.

In our case studies, we look for high structure similarity as a proxy for adopting correct function and low global sequence similarity to account for memorization. In Extended Data Figure 4c, we examine the Sinkhorn distance between function-conditioned generated latent embeddings and held-out real proteins with this GO term annotation, to examine distribution similarities between real and generated samples. To compare, we also examine the Sinkhorn distance between *random* proteins from the heldout set of real samples.

#### Generation and Filtering of Heme Proteins

For experimental validation, we focused on conditional generation of heme-binding proteins from the organism Escherichia coli (E. coli), which corresponded to function ID 17 and organism ID 1030. Although we additionally tested with unconditional organism generation, we did not notice a large difference in generated sequences based on *in silico* metrics, so we focus here on E. coli-conditioned generations.

We focused on two lengths for generation, 120 and 160 amino acids, respectively. For each length, we generated 1000 protein sequences and structures using the same configuration as *in silico* conditional evaluation and the PLAID model with 100 M parameters.

Starting from the two sets of 1000 generated sequences, we filtered down to those that were designable based on the structure prediction metrics used in *in silico evaluation*. From here, we additionally considered the sequences of length 120 that were inverse folded from PLAID-generated structures. Afterward, we aligned generated sequences to a known heme protein using mafft [61], and only retained those with a histidine located at the position necessary for coordinating heme. Using Foldseek, we filtered down to only sequences with a TM score *>* 0.5 compared to a known fold of a heme binding protein. We were then left with three sets of sequences, which are the “valid” sequences used for analysis in the main text. Those of length 120, length 160, and inverse folded of length 120 are referred to as Globin, H-NOX, and IF Globin proteins, respectively.

We then ran AlphaFold3 structure prediction to model the joint structure of the heme protein and the folded sequence; rather than searching for a multiple sequence alignment, we used the known structure of a heme-binding protein as the template. For the sequences of length 120 and 160, we filtered down to only structures with ipTM score greater than 0.7 and 0.6, respectively. The ipTM describes the inter-chain structure prediction confidence of the heme co-factor relative to the folded protein. For the non-inverse folded sequences, we only took those with a pLDDT *>* 60 as predicted by ESMFold, for the protein structure folded without heme. We then clustered the sequences at 90% sequence identity using mmseqs2 and took up to one sequence per cluster with the highest ipTM score. Maximum sequence identity to a known protein was calculated by running Diamond BLAST [62] against UniRef90 as a reference database.

### 6.9 Experimental Methods plasmid construction and transformation

The pET22b(+) vector was linearized by inverse PCR using Phusion polymerase (NEB, Ipswich, MA) and primers 007 and 008 (008 Forward: CTCGAGCACCAC-CACCACCACCACTGAGATCCGGC; 007 Reverse: CATATGTATATCTCCTTCT-TAAAGTTAAACAAAATTATTTC) digested with *DpnI* (NEB, Ipswich, MA), gel-purified, and validated to ensure minimal background transformation. The selected genes were synthesized by Twist Bioscience (South San Francisco, CA). DNA fragments containing flanking regions (upstream: GTTTAACTTTAAGAAGGAGATAT-ACAT; downstream: CTCGAGCACCACCATCACCACCACTGA) for Gibson assembly were received as dry residues in 96-well plates at up to 1 *µ*g per well. These DNA samples were dissolved in PCR-grade water to a concentration of approximately 50 ng/*µ*L. Following NEB recommendations, 1 *µ*L of DNA fragment was mixed with 1 *µ*L of linearized pET22b(+) (100 ng/*µ*L) and 5 *µ*L of Gibson assembly mix in a 96-well PCR plate. The plate was sealed and incubated at 50 °C for 60 min, then placed on ice. Subsequently, 5 *µ*L of chemically competent *E. coli* T7 Express cells (NEB, Ipswich, MA) were added to each well, followed by a 20 min incubation on ice and a 10 s heat shock in a 42 °C water bath. After transformation, 100 *µ*L of Luria–Bertani (LB) medium were added to each well, and 10 *µ*L of the mixture were used to inoculate 500 *µ*L of LB containing 100 *µ*g/mL ampicillin (LB_amp_) in a 96-deep-well plate and incubated at 37 °C for 16–18 h. Glycerol stocks were prepared for long-term storage.

#### UV-vis of all variants

Glycerol stocks were used to inoculate 300 *µ*L LB_amp_ in 96-well plates, covered with a sterile, breathable film, and grown at 37 °C overnight. From the stationary-phase cultures, 50 *µ*L were transferred into 900 *µ*L TB containing 100 *µ*g/mL ampicillin and supplemented with trace metal mix (TB_amp_) and incubated for 2 h at 37 °C prior to induction with with 50 *µ*L of IPTG in TB_amp_ (0.5 mM final). Induced cultures were incubated for 22 h at 22 °C to allow protein expression. Cells were then harvested by centrifugation (4,000×*g*, 5 min), and the resulting pellets were stored at −20 °C. For lysate preparation in 96-well plates, cell pellets were resuspended in 200 *µ*L of lysis buffer (100 mM KPi, pH 8.0, supplemented with 100 *µ*M PLP, 1 mg/mL lysozyme, 2 mM Mg^2+^, and DNase I) and incubated at 37 °C for 1 h with shaking at 200 rpm. Lysis was completed via three freeze–thaw cycles (≥5 min in an ethanol/dry ice bath, *>*30 min thaw at room temperature, followed by *>*5 min in a 37 °C water bath). Cell debris were removed by centrifugation (6,000×*g*, 15 min). In order to increase the concentration of proteins two plates were combined: 2 × 90 *µ*L clarified lysate from two plates were transferred into a transparent UV-Vis plate (Evergreen Scientific). The UV-Vis spectra of all variants were recorded using a Tecan Spark. The ferrous spectra were recorded by adding 20 uL of 300 mM dithionite.

#### Large-scale purification ad UV-vis

For large-scale expression, plasmids were isolated from selected colonies and sent for sequencing (Transnetyx, Inc., Cordova, TN). Valid plasmid were transformed into chemically competent *E. coli* T7 Express cells (NEB, Ipswich, MA) and single colonies were inoculated into LB_amp_ and grown overnight. The following day, 800 mL of TB_amp_ were inoculated at a 1:100 dilution from the overnight culture and incubated at 37 °C until the culture reached an OD_600_ of 0.6–0.8. Protein expression was induced with 0.5 mM IPTG and 1 mM ALA and allowed to proceed at 20 °C for 20–22 hours. Cells were harvested by centrifugation (5,000×*g*, 10 min) and washed once with PBS buffer (pH 7.4) and then stored at −20 °C until further use. For lysis, the cell pellet was thawed and resuspended in 50 mL lysis buffer (50 mM KPi, 200 mM NaCl, 20 mM imidazole, pH 8.0, DNAse I, lysozyme 1 mg/mL, 2 mM Mg^2+^) and incubated for 1 h at 37 °C. After incubation 50 *µ*M Hemin was supplemented to increase protein loading and lysed by sonication (3 min, 1 s on/2 s off, 35% amplitude). Lysates were clarified by centrifugation (*>*25,000×*g*, 45 min) and loaded onto 1-mL HisTrap columns using an AKTA Xpress system pre-equilibrated with buffer A (50 mM KPi, 200 mM NaCl, 20 mM imidazole, pH 8.0). Columns were washed with 10 column volumes (CV) of buffer A and proteins were eluted using a gradient to buffer B (50 mM KPi, 200 mM NaCl, 400 mM imidazole, pH 8.0). Eluted proteins were concentrated and buffer-exchanged with final buffer (50 mM KPi, 100 mM NaCl, pH 8.0) by centrifugation using Amicon Ultra-15 10 kDa MWCO (Merck Millipore Ltd.). Samples were used directly for UV-vis characterization. The purified proteins were normalized to a concentration of 20 *µ*M based on their predicted extinction coefficients at 280 nm and molecular weights. Spectra of the purified proteins were recorded using a Shimadzu UV-1800 spectrophotometer and a Starna quartz cuvette, corresponding to the oxidized (ferric) hemoprotein state. The cuvette was then degassed inside a Coy anaerobic chamber, after which 1 mM of freshly prepared dithionite was added. The cuvette was sealed, removed from the anaerobic chamber, and spectra were recorded immediately. Subsequently, CO gas was bubbled in the cuvette for 30 seconds, and a new spectrum was collected immediately after.

## 7 Declarations

### Data availability

Model weights for our 100M and 2B models are are available at https://huggingface.co/amyxlu/plaid/tree/main.

### Code availability

Code is publicly accessible at https://github.com/amyxlu/plaid.

### Author contributions

AXL designed the model with guidance from PA, WY, KKY and NF. AXL implemented and trained the models with guidance from WY. AXL performed in silico unconditional experiments with guidance from NF, and conditional sampling results with guidance from SR. JY performed sampling and filtering for heme binding experiments. YL and TL designed and performed wet-lab experiments experiments. AXL, JY, and TL wrote the initial draft. All authors contributed to the final draft. KKY, VG, KC, RB, NF, FA, and PA provided guidance throughout all stages.

## Footnotes

1 In practice, to make use of array broadcasting, a standard *M* is selected for all residues, with an associated one-hot mask to specify which atoms are present for a given residue. Following prior work [9, 52], we use the atom14 representation where *M* = 14.

## B.1 Defining the Latent Space: Pseudocode

To better clarify how we obtain the latent representation from ESMFold [9], we provide a code sketch for the relationship of this representation and the rest of the ESMFold architecture: Note that `self.esm s combine` and `self.esm s mlp` are both trained end-to-end with loss objectives from the original ESMFold paper.

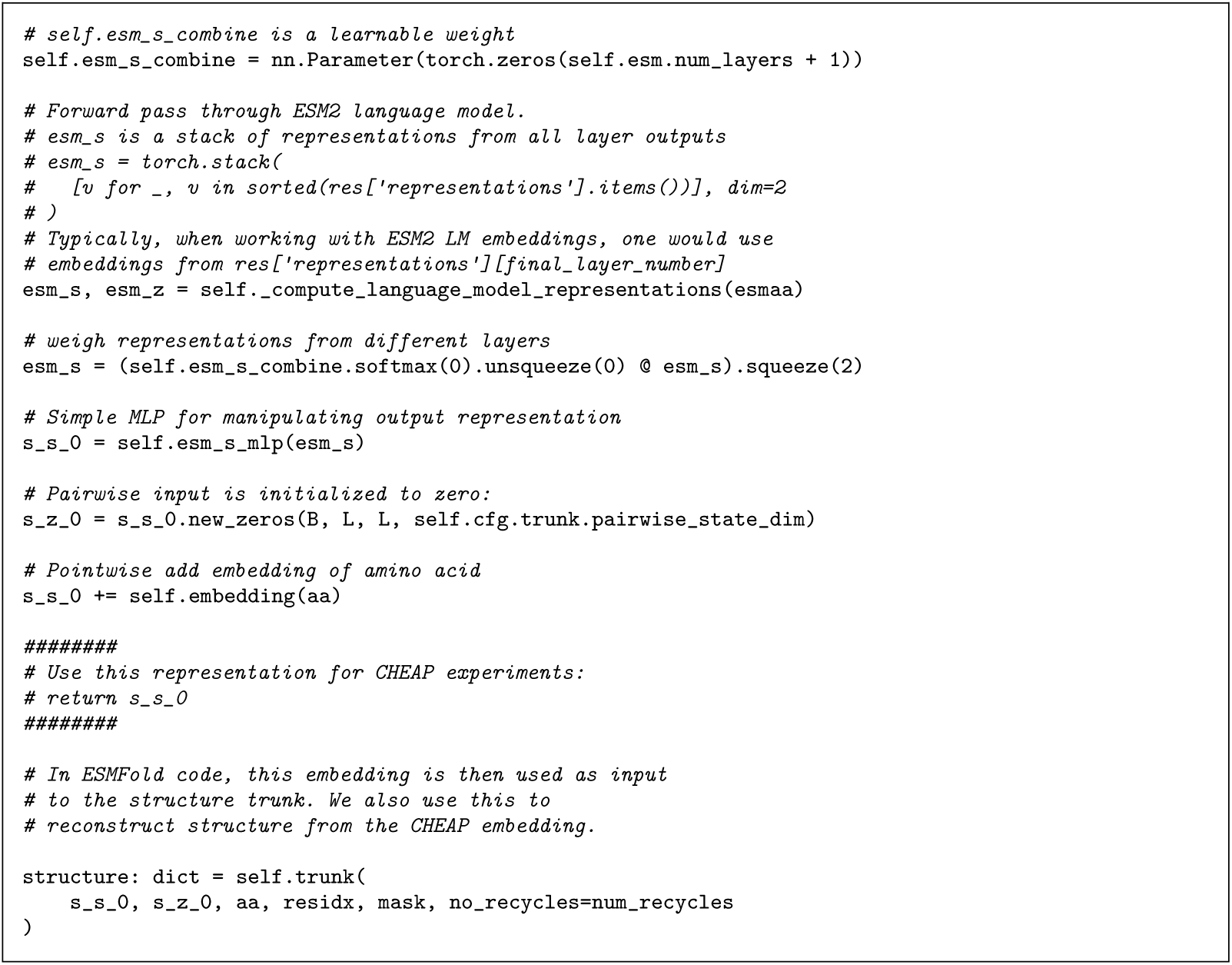

## B.2 CHEAP Compression Details

Briefly, the CHEAP encoder and decoder use an Hourglass Transformer [70] architecture that downsamples lengthwise, and downprojects the channel dimension, to create a bottleneck layer. The output of the encoder is our compressed embedding. The entire model is trained with the reconstruction loss MSE(x, ̂x). Results in Lu et al. [25] show that structural and sequence information in ESMFold latent spaces are in fact highly compressible, and despite using very small bottleneck dimensions, reconstruction performance can nonetheless be maintained when evaluated in sequence or structure space.

The embedding x, defined just before the structure module, is actually a linearly projected version of the ESM2 embeddings. If we defined x as the ESM2 embeddings directly, we could use the decoder from ESM2’s MLM training. However, since we use this modified embedding space, an approximation is necessary.

Based on reconstruction results in Lu et al. [25], we choose 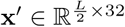 with L = 512, which balances reconstruction quality at a resolution comparable to the size of latent spaces in image diffusion models [58]. Dividing the length in half allows us to better leverage the scalability and performance of Transformers, while managing their O(L) memory needs.

The CHEAP module involves a channel normalization step prior to the forward pass through the autoencoder. We find that the distribution of embedding values is fairly “smooth” here (Figure ??). Althoughgh the original Rombach et al. [58] paper was trained with a KL constraint to a Gaussian distribution, we use the embedding output as is. CHEAP embeddings were also trained with a tanh layer at the output of the bottleneck; this allows us to clip our samples between [−1, 1] at each diffusion iteration, as was done in original image diffusion works [30, 59, 71, 72]. We found in early experiments that being able to clip the output values was very helpful for improving performance. Without using CHEAP compression prior to diffusion, sample quality was poor, even for short (L = 128) generations, as shown in Figure 3.

## B.3 Data

The PLAID paradigm can be applied to any sequence database. As of 2024, sequence-only databases range in size from UniRef90 [73] (193 million sequences) to metagenomic datasets such as BFD [74] (2.5 billion sequences) and OMG [75] (3.3 billion sequences). We use Pfam because it provides more annotations for in silico evaluation and because protein domains are the primary units of structure-mediated functions.

We use the September 2023 Pfam release, consisting of 57,595,205 sequences and 20,795 families. PLAID is fully compatible with larger sequence databases such as UniRef or BFD (roughly 2 billion sequences), which would offer even better coverage. We elect to use Pfam because sequence domains have more structure and functional labels, making it easier for in silico evaluation of generated samples. We also hold out about 15% of the data for validation.

We examine all Pfam domains that have a Gene Ontology mapping, resulting in 2,219 GO terms compatible with our model. For domains with multiple associated GO term labels, the GO term that is least prevalent in our dataset is selected, to encourage the selected terms to be more specific.

Approximately 46.7% of the dataset (N = 24, 637, 236) is annotated with a GO term. Using the publicly available mapping as of July 1, 2024, we count all GO occurrences; for each Pfam entry with multiple GO entries, we pick the one with the fewest GO occurrences to encourage more descriptive and distinct GO labels.

The Pfam-A.fasta file available from the Pfam FTP server includes the UniRef code of the source organism from which the Pfam domain is derived. The UniRef code furthermore includes a 5-letter “mnemonic” to denote the organism. We examine all unique organisms in our dataset and find 3,617 organisms. Samples are randomly cropped to a size of 512, which was selected by examining the distribution of all sequence lengths in Pfam (Extended Data Figure 4d).

## B.4 Sampling

Inference-time sampling hyperparameters provide the user with additional control over quality and sampling speed trade-offs. PLAID supports the DDPM sampler [30] and the DDIM sampler [71], as well as the improved speed samplers from DPM++ [67]. We find that using the DDIM sampler with 500 timesteps using either the sigmoid or cosine schedulers works best during inference, and reasonable samples can be obtained using the DPM++2M-SDE sampler with only 20 steps. Experiments shown here use the DDIM sampler with the sigmoid noise schedule at 500 timesteps.

Note that the performance bottleneck is found mostly during the latent sampling and structure decoding (which depends on the number of recycling iterations [9, 60] used); however, these two processes can be easily decoupled and parallelized, which cannot be done in existing protein diffusion methods. Furthermore, it allows us to prefilter which latents to decode using heuristic methods, and decode only those latents to structure, which would boost performance for nearly the same computational cost. We do not empirically explore this in this paper to provide a fair comparison, and because the filtering criteria would vary greatly by downstream use.

## B.5 Evaluation Details

For all benchmarks and models, we use default settings provided in their open-source code. For ProteinMPNN [5], we use the v 48 002 model with a sampling temperature of 0.1 and generate 8 sequences per protein, from which the best performing sequence is chosen. To calculate self-consistency, we fold sequences using OmegaFold [76] rather than ESMFold, again using default settings.

Though our models generate all-atom structure, we examine Cα RMSD rather than all-atom RMSD to avoid misattributing sequence generation underperformance to structure generation failures. Also, since there are usually differences in the sequences that are generated, different numbers of atoms make it difficult to assess all-atom RMSD.

For the hold-out natural reference dataset, we use sequences from Pfam and keep length distributions similar to that of the sampled proteins. Specifically, for each sequence bin between {64, 72, . . . , 504, 512}, we take 64 natural sequences of that length. For the experiment in Figure 4f, we use the Sinkhorn Distance rather than the Fŕechet Distance used commonly in images and video. Since not all functions have a large number of samples, we elected to use a metric that works better in low-sample settings.

Structure novelty is obtained by searching samples against PDB100 using Foldseek [36] easy-search. We examine the TM-score to the closest neighbor. For Foldseek and MMseqs experiments, all clustering experiments are performed by length. We use default settings for both tools. Though we report the average TM-Score to the top neighbor for Foldseek, we run easy-search in 3Di mode. For sequences, we use MMseqs2 [69] to see if sequences have a homolog in UniRef50, using default sensitivity settings. For samples with homologs, we further calculate the average sequence identity to the closest neighbor to assess novelty (Seq ID %).

## B.6 Distributional Conformity to Biophysical Attributes

For Wasserstein Distance to the distribution of biophysical attributes, we examine the following:

- Molecular Weight (MW): the molecular weight calculated from residue identities specified by the sequence.
- Aromaticity: relative frequencies of phenylalanine, tryptophan, and tyrosine, from Lobry and Gautier [77].
- Instability Index: dipeptide-based heuristic of protein half-life, from Guruprasad et al. [33].
- Isoelectricity (pI): the pH at which a molecule has no net electrical charge.
- Hydropathy: based on the GRand AVerage of hYdropathy (GRAVY) metric, from Kyte and Doolittle [34].
- Charge at pH = 7: the charge of a given protein at pH = 7, i.e., neutral pH.

---

**Supplemental Information Fig. 1:**
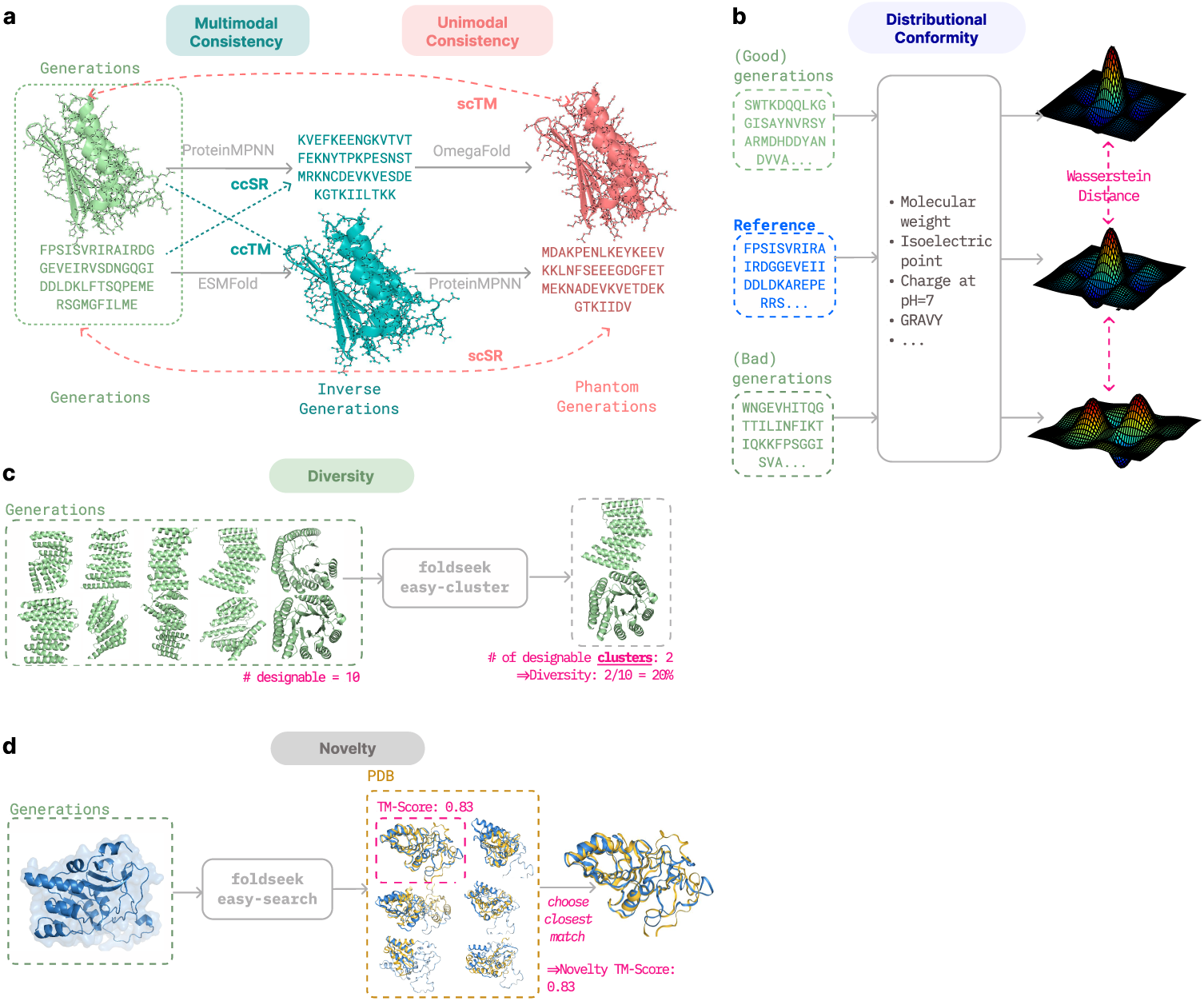
Schematic of metrics used to assess quality, diversity, and novelty. (a) Quality is assessed via multimodal and unimodal consistency metrics. (b) Distributional conformity is theWasserstein distance between biophysical attributes for generated proteins and a reference set of natural proteins. (c) Diversity is the number of unique designable (i.e. ccRMSD ! 2°A) samples. (d) Novelty is the TMScore between the generation and its most similar known protein. Sequence novelty and diversity are calculated similarly, but are omitted from the schematic for simplicity: we use mmseqs easy-search and mmseqs easy-cluster [69] instead of foldseek [36] to do sequence-level search/clustering, and novelty similarity is calculated as the sequence identity after pairwise alignment.

**Supplemental Information Fig. 2:**
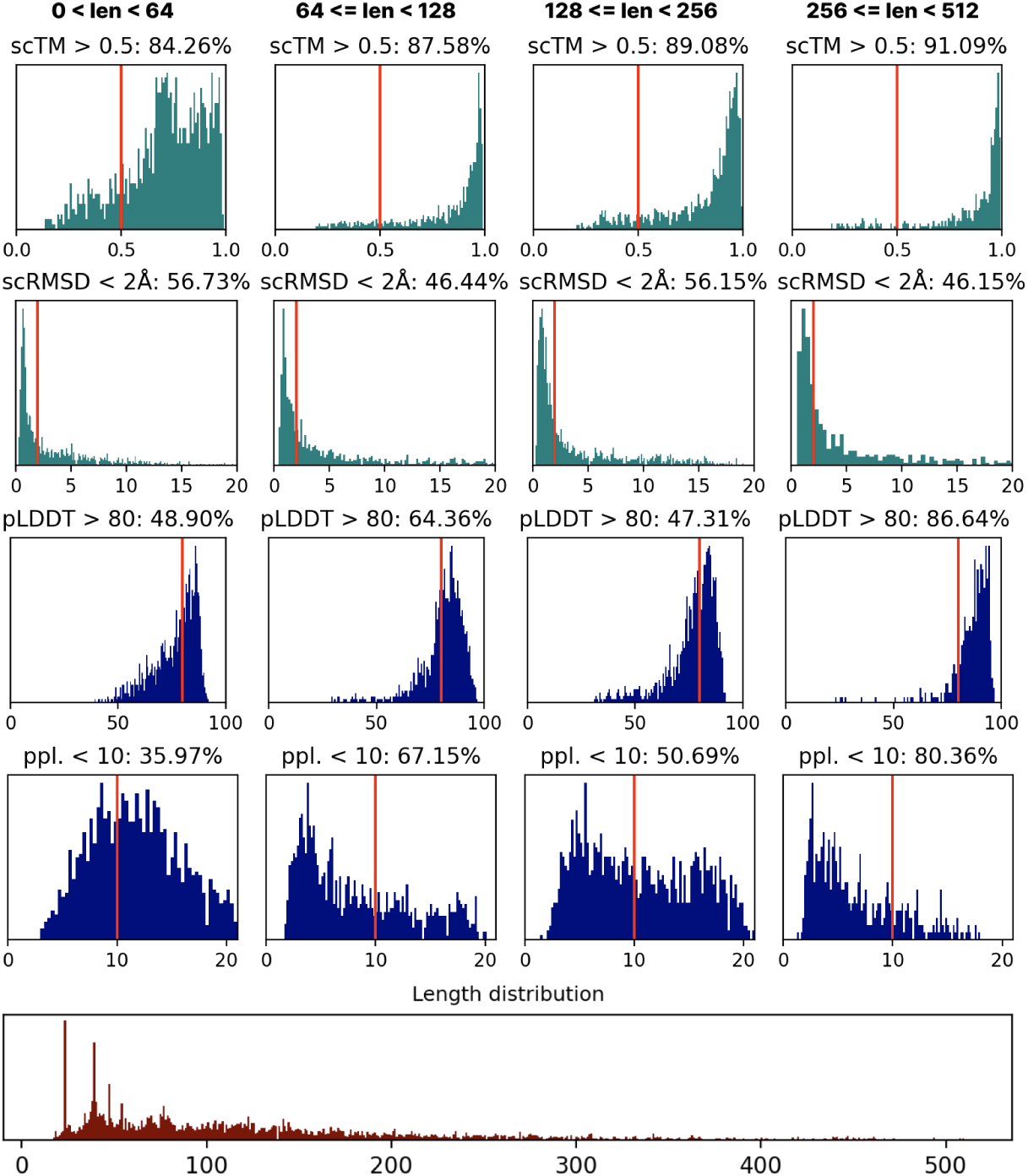
In order to examine the usefulness of in silico quality metrics, we plot scTM, scRMSD, pLDDT, and perplexity for natural proteins. As shown, it is not always useful to aim for strict optimization of these metrics, since even natural proteins do not always have an idealized value for these metrics.

**Extended Data Fig. 1:**
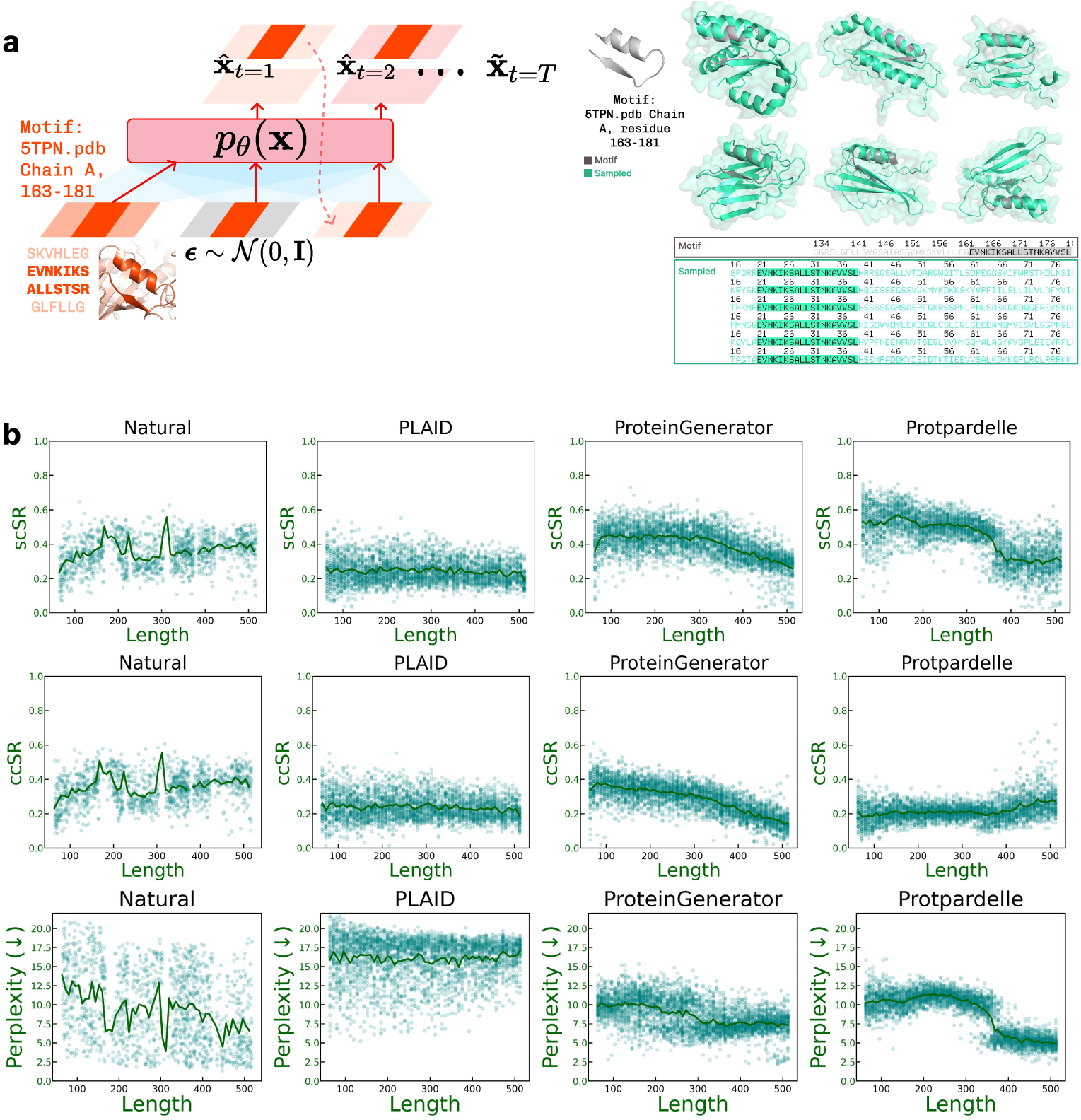
Motif conditioning and length-stratified sequence analysis. (a) Schematic of motif-constrained generation. A fixed structural motif (PDB: 5TPN, chain A, residues 163-181) is embedded into the latent trajectory, and PLAID is conditioned to preserve its coordinates while freely generating the surrounding sequence and structure. Representative sampled structures (right) show consistent placement of the motif (green) across diverse scaffolds, and corresponding sequence logos illustrate recovery of motif-proximal residues. (b Examining sequence-related metrics for co-generation methods. scSR (self-consistency sequence recovery), ccSR (cross-consistency sequence recovery), and perplexity metric calculations are detailed in Supplemental Information Figure 1. As noted in Figure 3c, at longer lengths, methods may be prone to generating many repeats, which is difficult to catch using perplexity and recovery-based metrics.

**Extended Data Fig. 2.**
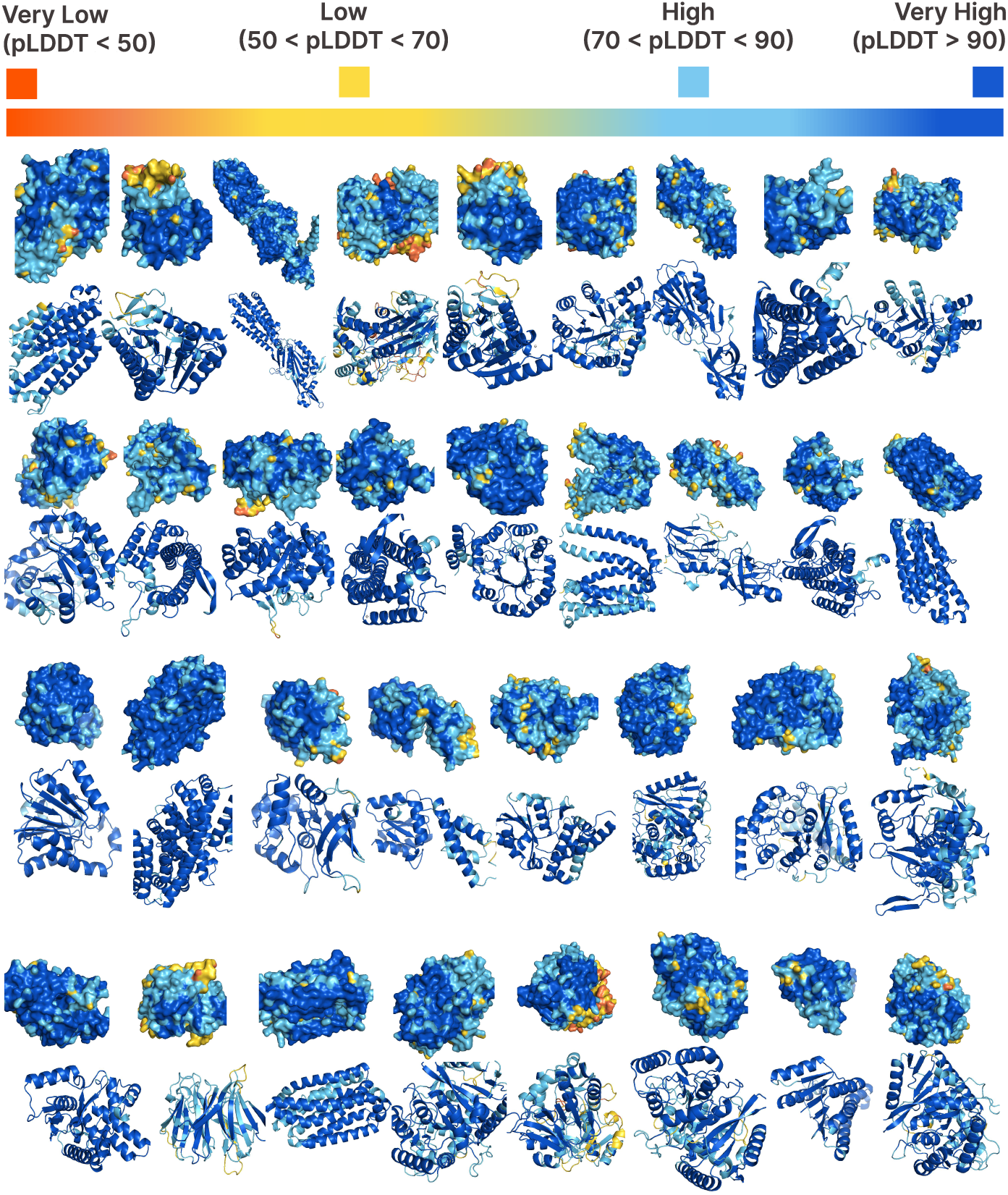
PLAID unconditionally generates diverse, high-quality allatom structures, despite using only sequences for training the generative model.

**Extended Data Fig. 3:**
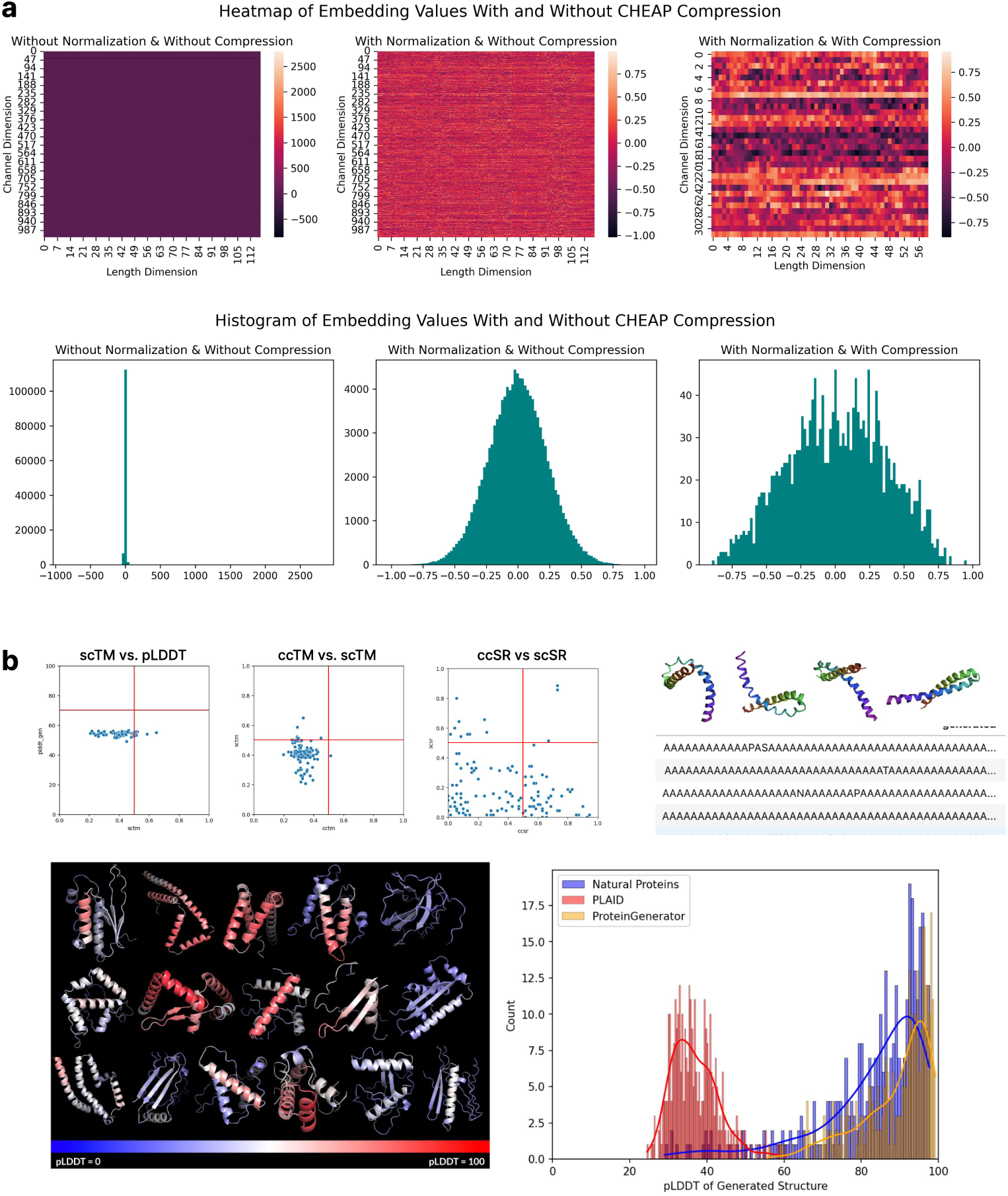
(a) Visualizing the original ESMFold latent space before normalization, after per-channel normalization, and after compression. Without compression, values are unsmooth. After compression, the value distribution is smoother and more amenable to being learned by a generative model. (b) Results when running PLAID on the ESMFold latent space naively without CHEAP compression, for proteins of length 128. There is a tendency to generate repeated sequences, and quality is low compared to baselines.

**Extended Data Fig. 4:**
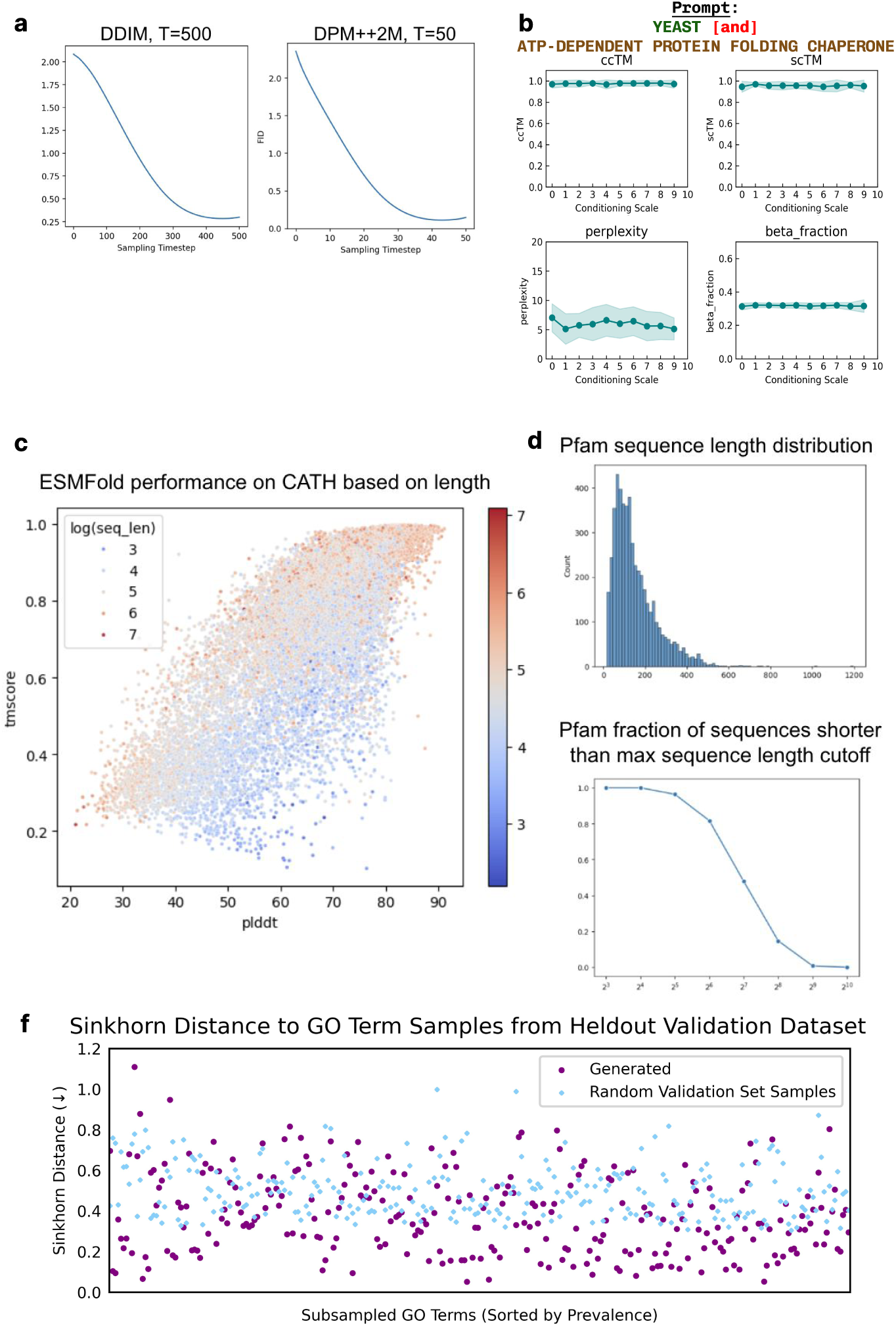
(a) PLAID supports using DPM++2M [67], which allows for diffusion sampling with much fewer sampling steps. Here we show results using the DPM++2M sampler using 20 steps, versus the default PLAID sampling (500 steps using the DDIM [31] sampler. The y-axis is the Frechet distance between the generated embedding and a reference set of true embeddings (lower is better). We can achieve similar results of a similar quality using the DPM++2M sampler. (b) Examining the effect of conditioning weight on generation quality. (c) Understanding length generalization for ESMFold; this acts as sanity check for understanding the protein lengths at which the ESMFold structure decoder can produce good results, and to understand whether if pLDDT is useful as an evaluation metric. We run ESMFold over the CATH [68] dataset and assess the pLDDT (model confidence) against the prediction accuracy (TM-score). Defining model hallucination to be a case when a model is confident, but wrong, we find this to be much more common in shorter sequences. (d) Understanding which length cutoff to use during training. We use a cut off of 512 based on the findings here, since 512 would allow us to see full domains for most sequences in the dataset, while still having manageable GPU memory needs. (f) Examining the Sinkhorn distance between GO-term conditioned samples and a collection of true samples with the same GO term, at the embedding generation level. For reference, we also compare it to the Sinkhorn distance between a random set of real proteins from the validation set, and to the GO-term specific collection of real proteins.

**Extended Data Table 1:** Comparison of model performance across consistency and quality metrics. Arrows indicate whether higher (↑) or lower (↓) values are better. Bold values show best performance among all-atom generation models. pLDDT refers to the confidence score directly returned by the structure trunk of the generative model; for models which do not produce a pLDDT metric, N/A is used.

|  | Cross-Modal Consistency |  |  |  | Structure Quality |  |  | Sequence Quality |  |
| --- | --- | --- | --- | --- | --- | --- | --- | --- | --- |
| | ccTM (↑) | ccRMSD (↓) | ccSR (↑) | % ccRMSD < 2Å (↑) | scTM (↑) | pLDDT (↑) | Beta sheet % (↑) | scSR (↑) | Ppl. (↓) |
| ProteinGenerator | 0.58 | 11.86 | **0.28** | 0.08 | **0.72** | **69.00** | 0.04 | 0.40 | **8.60** |
| Protpardelle | 0.44 | 24.28 | 0.22 | 0.00 | 0.57 | N/A | 0.11 | **0.44** | 8.86 |
| PLAID | **0.69** | **9.47** | 0.26 | **0.32** | 0.64 | 59.46 | **0.13** | 0.27 | 14.61 |
| *Natural* | *1.00* | *0.07* | *0.39* | *1.00* | *0.84* | 84.51 | 0.13 | 0.39 | 7.40 |

**Extended Data Table 2:**
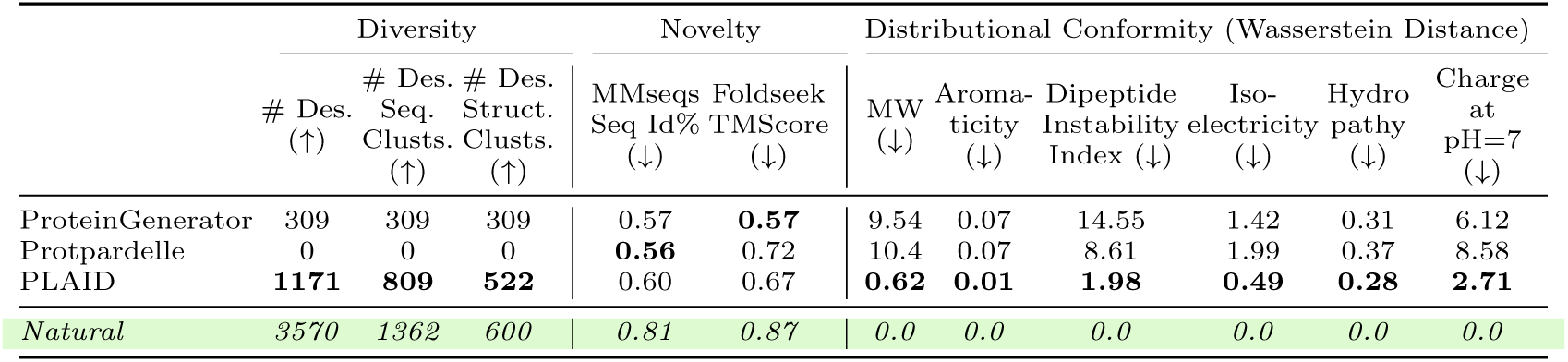
Diversity, novelty, and distributional conformity. metrics across models. Bold values show best performance among all-atom generation models. Descriptions of each biophysical parameter for distributional conformity is described in Appendix B.6.

**Extended Data Table 3:** Ablation results.

| Config | Configuration | ccTM | scTM | Ppl. | Seq. Div. % | Struct. Div. % |
| --- | --- | --- | --- | --- | --- | --- |
| A | cosine noise sched. & pred. noise | 0.54 | 0.55 | 16.97 | **0.98** | 0.86 |
| B | A + v-diffusion | 0.52 | 0.53 | 17.37 | **0.98** | **0.89** |
| C | A + MinSNR | 0.59 | 0.59 | 16.76 | 0.97 | 0.86 |
| D | A + B + C + sigmoid noise sched. | 0.56 | 0.58 | 16.88 | 0.92 | 0.86 |
| **E** | **D + self-conditioning** | **0.70** | **0.65** | **15.38** | 0.93 | 0.76 |
| F | E + no cond drop | 0.57 | 0.57 | 17.28 | 0.97 | 0.85 |

**Extended Data Table 4:** Time required to sample proteins with 600 residues. . We assess time required both for sampling *N* = 100 samples. Experiments are run on Nvidia A100. Methods marked by (*) do not support batching in the default implementation; we assess both batched and unbatched runtimes. A rate limiting step for PLAID is the structure decoder step; a pipeline implementation would allow both the latent sampling step and decoding to happen concurrently.

|  | seconds/sample, batched |  | seconds/sample, unbatched |  |
| --- | --- | --- | --- | --- |
|  | Sample Latent | Decode | Sample Latent | Decode |
| Protpardelle | 11.21 | - | 17.16 | - |
| Multiflow* | 231.32 | - | 277.11 | - |
| ProteinGenerator* | 343.32 | - | 342.28 | - |
| **PLAID (100M)** | 1.64 | 15.12 | 27.63 | 1.07 |
| **PLAID (2B)** | 19.34 | 15.07 | 49.03 | 0.9 |

**Extended Data Table 5:** Forward pass benchmark of vanilla multihead attention compared to the optimized xFormers implementation of memory-efficient attention [64] and FlashAttention-2 [65]. Though FlashAttention2 performed best in our benchmarks, a fused kernel implementation with key padding was not yet available at the time of writing. Since our data contained different lengths (as compared to most image diffusion use cases, or language use-cases that can make use of the implemented causal masking), we instead use the xFormers implementation. We expect that sampling speed results would improve once this feature is becomes available in the FlashAttention package.

| Method | Mean Time (s) | Mean Memory (GB) |
| --- | --- | --- |
| Standard Multihead Attention | $0.0946 \pm 9.23\text{e-}4$ | $76.0 \pm 0.409$ |
| xFormers Memory Efficient Attention | $0.0519 \pm 4.33\text{e-}05$ | $64.0 \pm 0.409$ |
| Flash Attention | $0.0377 \pm 1.91\text{e-}3$ | $49.2 \pm 0.783$ |

**Extended Data Table 6:**
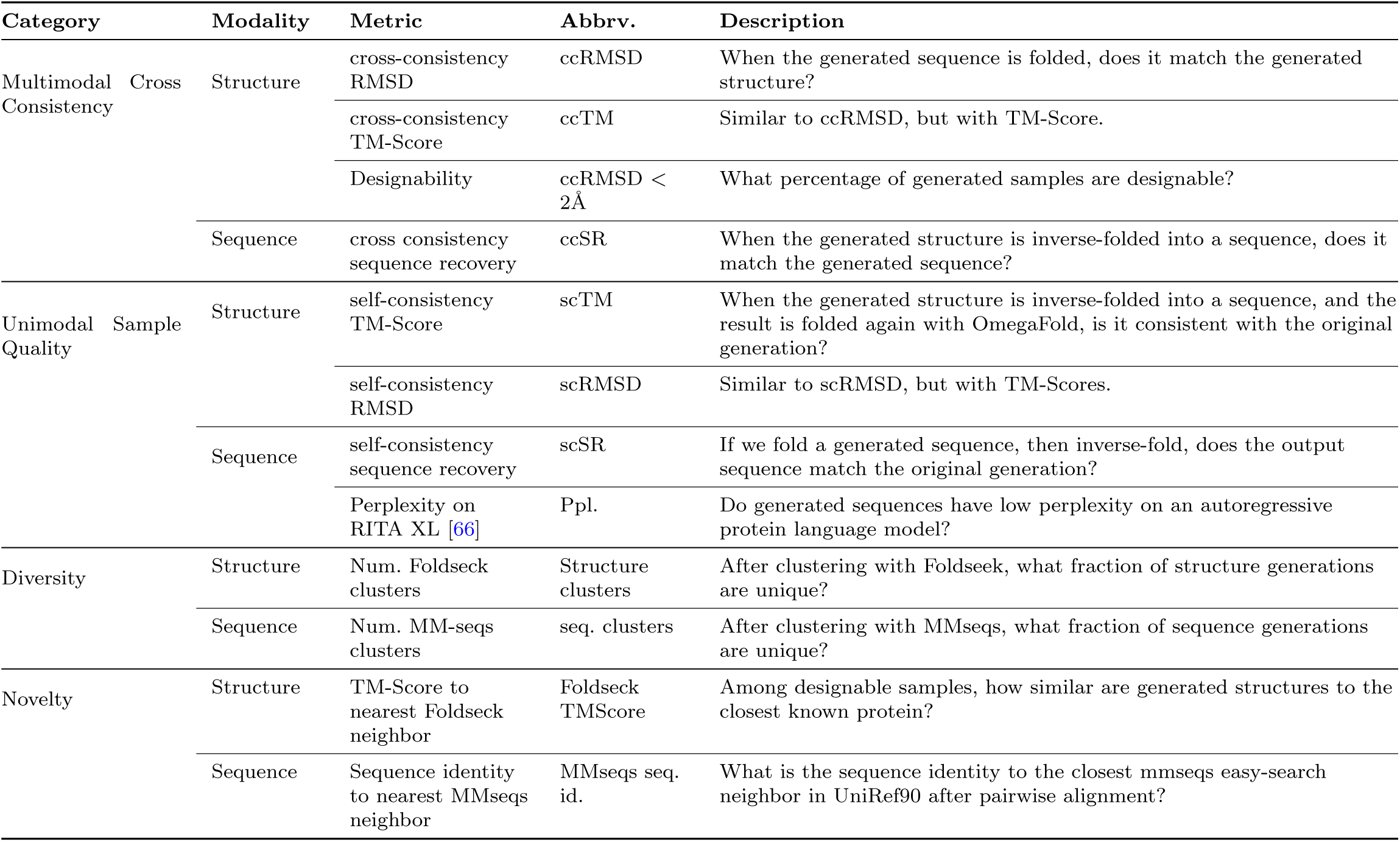
Evaluation metrics for protein structure and sequence generation.

